# Genipin Crosslinks the Extracellular Matrix to Rescue Developmental and Degenerative Defects, and Accelerates Regeneration of Peripheral Neurons

**DOI:** 10.1101/2023.03.22.533831

**Authors:** Kenyi Saito-Diaz, Paula Dietrich, Hsueh-Fu Wu, Xin Sun, Archie Jayesh Patel, Camryn Gale Wzientek, Anthony Robert Prudden, Geert-Jan Boons, Shuibing Chen, Lorenz Studer, Bingqian Xu, Ioannis Dragatsis, Nadja Zeltner

## Abstract

The peripheral nervous system (PNS) is essential for proper body function. A high percentage of the population suffer nerve degeneration or peripheral damage. For example, over 40% of patients with diabetes or undergoing chemotherapy develop peripheral neuropathies. Despite this, there are major gaps in the knowledge of human PNS development and therefore, there are no available treatments. Familial Dysautonomia (FD) is a devastating disorder that specifically affects the PNS making it an ideal model to study PNS dysfunction. FD is caused by a homozygous point mutation in *ELP1* leading to developmental and degenerative defects in the sensory and autonomic lineages. We previously employed human pluripotent stem cells (hPSCs) to show that peripheral sensory neurons (SNs) are not generated efficiently and degenerate over time in FD. Here, we conducted a chemical screen to identify compounds able to rescue this SN differentiation inefficiency. We identified that genipin, a compound prescribed in Traditional Chinese Medicine for neurodegenerative disorders, restores neural crest and SN development in FD, both in the hPSC model and in a FD mouse model. Additionally, genipin prevented FD neuronal degeneration, suggesting that it could be offered to patients suffering from PNS neurodegenerative disorders. We found that genipin crosslinks the extracellular matrix, increases the stiffness of the ECM, reorganizes the actin cytoskeleton, and promotes transcription of YAP-dependent genes. Finally, we show that genipin enhances axon regeneration in an *in vitro* axotomy model in healthy sensory and sympathetic neurons (part of the PNS) and in prefrontal cortical neurons (part of the central nervous system, CNS). Our results suggest genipin can be used as a promising drug candidate for treatment of neurodevelopmental and neurodegenerative diseases, and as a enhancer of neuronal regeneration.

**One sentence summary:** Genipin rescues the developmental and degenerative phenotypes of the peripheral neuropathy familial dysautonomia and enhances neuron regeneration after injury.

## Introduction

The peripheral nervous system (PNS) consists of sensory, sympathetic, parasympathetic, and enteric neurons, all of which develop from the neural crest (NC). The PNS innervates all organs of the body and is essential for its homeostasis. Loss, as well as hyper-function of the PNS lead to an array of human disorders caused by genetic mutations, metabolic problems, traumatic injury, inflammation, toxins, and infections. While lots of research is invested in finding treatments^1^, to date, there is no FDA approved treatment available. For example, peripheral nerve degeneration (peripheral neuropathy) and peripheral nerve damage are common, i.e. 20 million people and 20-30% of the population, respectively are sufferers. Together, they cause a large impact on life and are a major healthcare cost burden^2, 3^.

To improve our general understanding of PNS function and missregulation, it is useful to study a model disease; ideally one that is well defined, specific and causes a strong phenotype^4^. The autosomal recessive childhood disorder Familial Dysautonomia (FD) serves as such a model disorder. FD is caused by a homozygous point mutation in the gene *ELP1* (formerly *IKBKAP*), the scaffolding component of the transcriptional elongator complex^5^. The mutation leads to aberrant splicing and tissue-specific reduction of functional ELP1 protein^6^. This reduction is particularly prominent in the neural crest (NC) progenitor lineage and its progeny: the peripheral nervous system (PNS), especially in sensory and autonomic neurons^6^. These neurons develop at a reduced rate and they degenerate rapidly over time^7^. As a result, patients experience loss of pain perception, ataxic gait, trouble regulating heart rate and blood pressure and debilitating dysautonomic crisis, characterized by tachycardia, blood pressure spikes and extensive vomiting^2, 8^. Although FD mouse models show the same phenotypes: reduced size and neuron numbers in the dorsal root ganglia (DRG)^9^, the mechanism of how SNs fail to develop and quickly degenerate remains elusive. Thus, none of the interventions available to FD patients target SN symptoms^10^. However, this problem is not limited to FD: other patients with SN loss or defects stemming, for example from peripheral neuropathies or cancer chemotherapy treatment have limited pharmacological options as well^11^. Thus, a drug that targets SNs is necessary to treat a wide range of diseases.

Human pluripotent stem cell (hPSC) technology^12^ is ideal to gain cellular and mechanistic insight into genetic, early-onset human disorders; as it allows the retrieval of patient specific cells, their reprogramming into hPSCs, followed by their differentiation into SNs^13^, and parallel comparison between healthy control and disease cells for disease mechanistic studies^14^. Additionally, it provides the possibility to generate large numbers of neurons that can be employed for high-throughput drug screening approaches. In fact, FD was one of the first diseases that was modeled using the hPSCs technology^15^ and provided one of the first proof-of-principle drug screens employing hPSCs^16^. We previously showed that this disease modeling technique is highly sensitive in that it allows the recapitulation of specific phenotypes accounting for patient’s varying severity of symptoms^17^. We showed that in FD, both NC cells and SNs are not developing efficiently and that the SNs degenerate/die over time *in vitro*. These findings are consistent with patients being born with less SNs in the DRG^7^, equivalent findings in the FD mouse DRG^9^ and the fact that patient’s pain/temperature perception decreases with age^18^.

Here, we conducted a high-throughput chemical screen and identified genipin as a hit compound that targets SN defects in FD. Genipin is derived from the fruit of *Gardenia Jasminoides* and is the active ingredient of the Traditional Chinese Medicine (TCM) Zhi Zi^19^, which is being prescribed for neurodegenerative disorders, including Alzheimer’s^20, 21^ and Parkinson’s disease^22, 23^. In this context genipin has shown a safe profile in humans. Genipin has been implicated as an inhibitor of UCP2, preventing UCP2-mediated proton leak in mitochondria and increasing mitochondrial membrane potential^24^. Furthermore, genipin has been shown to have anti-inflammatory effects by inhibiting nitric oxide synthesis *in vitro* and *in vivo*^25^. Lastly, genipin is being used by bioengineers to crosslink extracellular matrices (ECM), due to its low toxicity^26, 27^. We show that genipin reverses the differentiation inefficiency observed in FD NC and SNs and prevent degeneration of SNs *in vitro*. We also demonstrate that genipin can be fed to pregnant FD mice to rescue the neurodevelopmental phenotypes in embryos. We provide evidence that genipin acts via ECM crosslinking and reverses FD phenotypes by activating the transcription of YAP-dependent genes. Lastly, we show that genipin promotes rapid regeneration of neurons from the peripheral (sensory and sympathetic) and central (prefrontal cortex) nervous systems. Together, our data provides support that genipin is a novel drug candidate to treat neurodevelopmental diseases, prevent neurodegeneration, and promote neuronal regeneration upon injury. Thus, genipin has the potential to be used as a treatment for a broad range of neurological diseases.

## Results

### Chemical screen to rescue sensory neuron phenotypes in Familial Dysautonomia

We first reproduced our previous findings that hPSC-derived SNs from FD patients cannot be generated efficiently *in vitro*^17^. The differentiation protocol defined by Chambers et al.^28^ (**Fig. 1A**) was employed to differentiate hPSCs, derived from healthy control embryonic stem cells (H9/WA09, hPSC-ctr-H9), healthy control iPSCs (iPSC-ctr-C1) and two iPSC lines from FD patients (iPSC-FD-S2 and iPSC-FD-S3)^17^. While the undifferentiated colonies looked normal in all four lines, once SNs were generated by day 13 of the differentiation protocol, the efficiency/number of neurons dropped dramatically in the FD lines compared to the control lines (**Fig. 1B**). Based on this phenotype, we set out to conduct a chemical drug screen to identify compounds that could reverse this phenotype and allow FD-iPSCs to efficiently generate SNs. We first adapted the SN differentiation protocol to high-throughput screening conditions. We tested the following adaptation parameters: 96 and 384 well plate formats, feeding frequency of twice (day 2, 5) or three times (day 2, 5, 8; normal, non-screening feeding is done every other day), shortening the protocol to 10 days total and thus removing the replating step at day 12, and replacing the media gradient with a 1:1 mixture of the KSR and N2 media, respectively instead of the gradual change of media (**Fig. 1C**). The simplifications of the protocol lead to an overall decrease in SN numbers from ∼60% (in the regular protocol^28^ to ∼30% (in our screening protocol) in hPSC-ctr-H9 cells. Within these conditions, the 96-well format and 3 times feeding produced the most robust SNs in the healthy control (**Fig. 1D**) and the largest difference between control and FD (**Sup. Fig. 1A, B**). We used these conditions to conduct a pilot screening using only positive (hPSC-ctr-H9) and negative (iPSC-FD-S2) controls. In these conditions, the calculated Z’ factor for the screen was 0.14, which denotes a marginal assay. However, thanks to the design and read-out of our screening, positive and negative controls were clearly distinguishable. We planned for a relatively small screen and thus proceeded with confidence. We reasoned that treatment starting on differentiation day 2 would allow initial specification into ectoderm but influence differentiation into NC and SN fates. We then screened 640 compounds, i.e. half of the LOPAC chemical library <u>o</u>f <u>p</u>harmacological <u>a</u>ctive <u>c</u>ompounds from Sigma, that contains a mixture of compounds from the fields of cell signaling and neuroscience (**Fig. 1E**). Each compound was screened at 1 mM and 10 mM in DMSO. All wells were then stained for the pan-SN marker *BRN3A* and DAPI and hit compounds were called if the fold change (FC) over the average of all DMSO wells was above the average of all compounds plus 3 SDs (**Fig. 1F, Sup. Table 1, 2**). Our screen resulted in 3 hits: Fluphenazine dihydrochloride (Flu, at 10 mM, a dopamine receptor antagonist), AC-93253 iodide (AC, at 1 mM, a retinoic acid receptor agonist) and genipin (at 1 mM and 10 mM, **Sup. Fig. 1C**). The controls on the screen were: Healthy hPSC-ctr-H9 and iPSC-FD-S2, both in DMSO, and DMSO alone. The Z-score for genipin was 15 at 1 mM and 17 at 10 mM.

**Figure 1.**
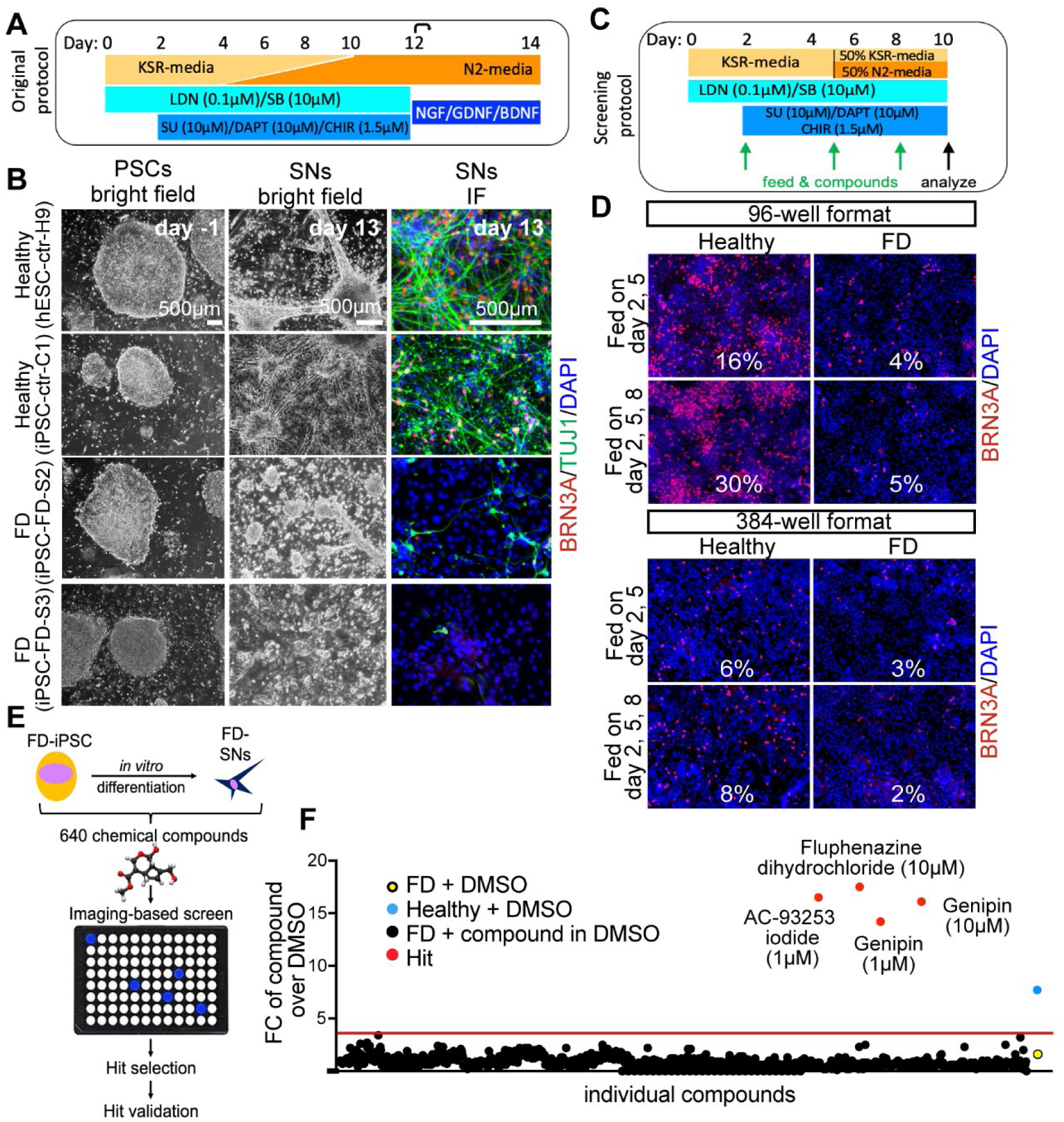
Chemical screen on FD sensory neurons. **A)** Differentiation protocol adapted from Chambers et al., 2012. **B)** Bright filed imaging shows normal morphology of undifferentiated hPSCs in all lines (left column). SN differentiation is impaired in both FD lines, but normal in healthy control lines (right two columns). **C)** Differentiation protocol adapted to high-throughput screening conditions. **D)** The SN differentiation protocol is most efficient in 96 wells when the cells are fed three times. **E)** Cartoon of the screen set up. **F)** 640 compounds from the LOPAC chemical library (the first half) were screened. Controls included DMSO only wells, healthy hPSC-ctr-H9 with DMSO (blue dot = positive control), iPSC-FD-S2 iPSCs with DMSO (yellow dot=negative control). Hit compounds were called when the fold change (FC) over the average of all DMSO wells was above the average of all compounds plus 3 SDs (here 3.8, red dots). Every compound was screened at 1 mM and 10 mM. 16 images were taken for each well, all wells were imaged for the ratio of BRN3A^+^/DAPI^+^ staining.

### Hit validation

Next, we performed several validation assays to further assess the hit compounds. The 3 hit compounds were tested in a repeat of the same conditions as performed in the screen itself (**Sup. Fig. 1D**) and then repeated in a different well format (48-wells) using the non-screening differentiation protocol (**Sup. Fig. 1E**). This analysis revealed that indeed genipin rescues the SN defect in FD (**Sup. Fig. 1E**, red rectangle), whereas Flu did not have much of a rescue effect and was auto-fluorescent (**Sup. Fig. 1E**, blue rectangle), and AC did not rescue the FD phenotype and showed high cytotoxicity (DAPI staining, **Sup. Fig. 1E**, green rectangle). We also tested the three hits on additional cell lines, including healthy iPSC-ctr-C1 and a FD line from patients with milder symptoms (iPSC-FD-M2)^17^. We observed the same phenotypes in iPSC-ctr-C1 and iPSC-FD-M2 cells (**Sup. Fig. 1F**). Thus, we focused on genipin. We titrated the genipin concentration (**Fig. 2A, B**) and found that, while genipin increased the number of SNs in a dose dependent manner in control hPSC-ctr-H9 and iPSC-ctr-C1cells (about 7-fold), the increase of SNs in genipin treated iPSC-FD-S2 cells was dramatically larger (over 30-fold). To further confirm genipin’s effect, we tested it in our newer SN differentiation protocol that is fully chemically defined (E8/E6)^13^ (**Sup. Fig. 1G**). Indeed, we found that genipin robustly rescued both the NC and the SN defect in iPSC-FD-S2 cells (**Fig. 2C**). However, in this protocol the cells were more sensitive, showing mild cytotoxic effects at 20 µM (likely due to the lack of the buffering KSR), thus the ideal concentration of genipin was lower (10 µM) (**Fig. 2C-E**). All data from here on was done in the chemically defined SN protocol^13^. Lastly, we tested genipin from different commercial sources (Sigma and Biomaterials) and found that, while both compounds worked, genipin from Biomaterials showed a more robust phenotype at 10 µM. Further experiments were done using genipin from this source (**Fig. 2F**). Together, we show that our small molecule screen resulted in the identification of genipin as a rescue compound in our FD model.

**Figure 2.**
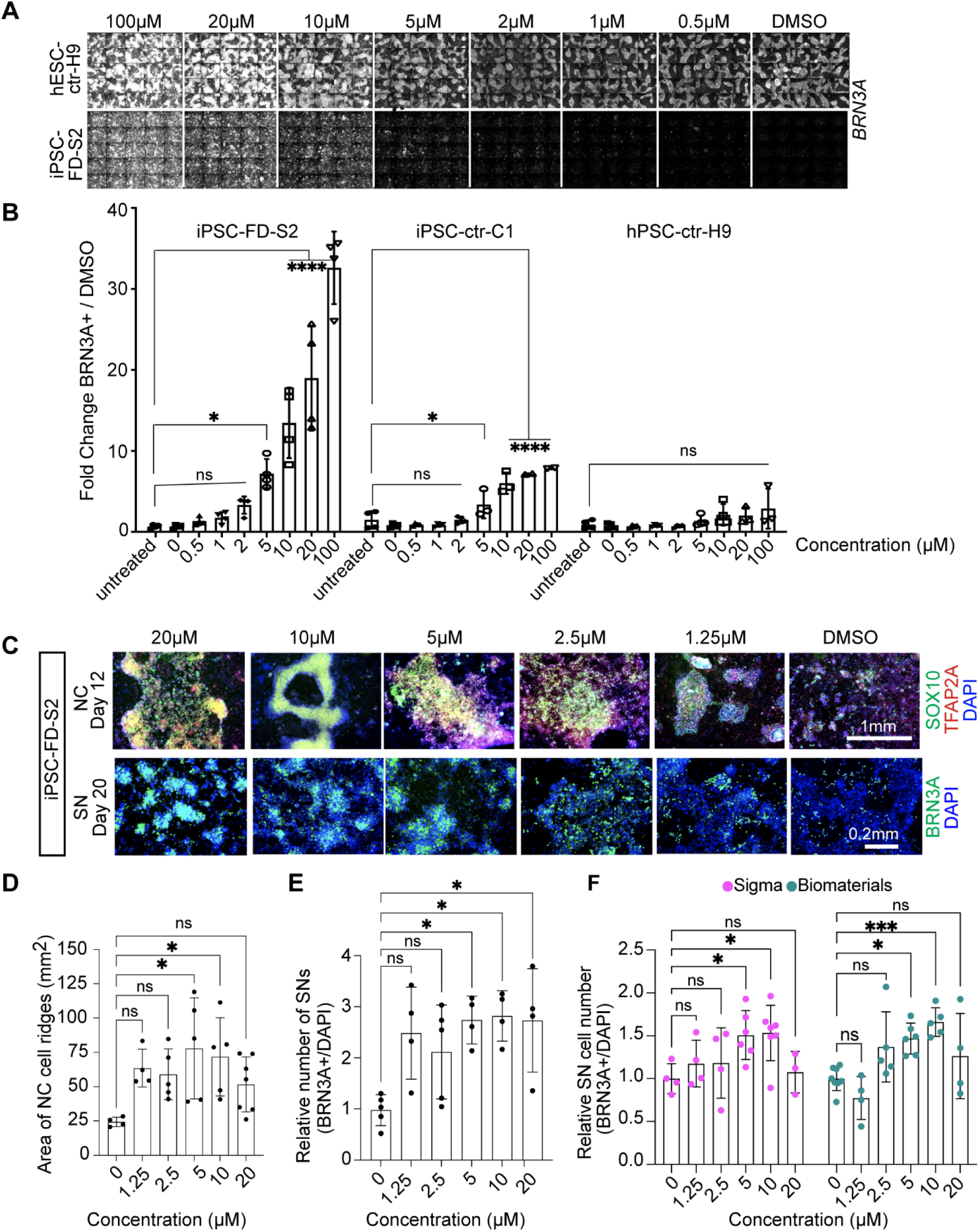
Validation of the hit compound genipin. **A)** Titration of genipin during the SN differentiation in healthy hPSC-ctr-H9 and iPSC-FD-S2 iPSCs. Differentiation protocol depicted in Fig. 1a was used. **B)** Quantification of genipin titration based on IF of BRN3A^+^ SNs. n=3-4 biological replicates. **C)** Titration of genipin in iPSC-FD-S2 cells during differentiation into SN-biased neural crest cells (top row) and SNs (bottom row). Cells were treated with indicated concentrations of genipin starting on day 2. Cells were then fixed on the indicated days and stained for SOX10 and TFAP2A (top) or BRN3A (bottom) and DAPI. **D and E)** Quantification of size of NC ridges and number of SNs upon genipin treatment. **D)** Area of ridges in **c** marked by DAPI staining (n=4-7 biological replicates) and **E)** number of BRN3A^+^ SNs in **c** were quantified (n=3-5 biological replicates). **F)** Genipin commercially obtained from both Sigma and Biomaterials rescues SN differentiation in iPSC-FD-S2. Cells were differentiated in the presence of genipin from either source starting on day 2. Cells were fixed on day 20 and stained for BRN3A and DAPI. n=3-7 biological replicates. All graphs show mean ± s.d. For **B**, **D**, **E**, and **F,** one-way ANOVA, *p<0.05, **p<0.005.

### Effects of genipin on neurodevelopment of neural crest cells and sensory neurons in FD iPSCs

We sought to carefully characterize genipin’s effects throughout development. In human embryonic development, NC cells (marked by *SOX10*) give rise to SN-specified NC cells (marked by *NGN1/2*), followed by *BRN3A*^+^ SNs that also express *ISL1*, *PRPH*, *TRKA*, *TRKB*, and *TRKC*. Thus, we first assessed the effects of genipin on NC formation. We found an increase of typical dark NC ridges that correlate with *SOX10* expression (on mRNA and protein level) in all the cell lines tested (**Fig. 3A-C**, arrows and **Sup. Fig. 2A**). Additional NC genes (*FOXS1* and *P75NTR*) were restored by genipin (**Sup. Fig. 2B**). We found that genipin increased *TFAP2A*, however non-significant, probably because *TFAP2A* is also expressed in non-neural ectoderm (**Sup. Fig. 2B**). We then confirmed that genipin further is capable of rescuing SN development by increasing the number of SNs in FD via staining for *BRN3A*, *TUJ1*, *ISL1*, and *PRPH* (**Fig. 3D, Sup. Fig 2C**) and quantification of the number of BRN3A^+^ positive neurons via intracellular FACS analysis (**Fig. 3E, Sup. Fig. 2D**). Furthermore, the expression of various SN genes (*BRN3A*, *ISL1*, *PRPH*, *TRKA*, *TRKB*, and *TRKC*) was rescued in FD by genipin, shown by RT-qPCR (**Fig. 3F, Sup. Fig. 2E**) and immunoblotting (**Fig. 3G**). Finally, we show that the firing rate in FD SNs also increases upon treatment with genipin compared to DMSO (**Fig. 3H**), likely because of the increased number of differentiated SNs. From here on forward, we pooled iPSC-FD-S2 and iPSC-FD-S3 data, also indicated in the figure legends. In sum, our results show that genipin rescues FD neurodevelopmental defects at the NC and SN stage.

**Figure 3.**
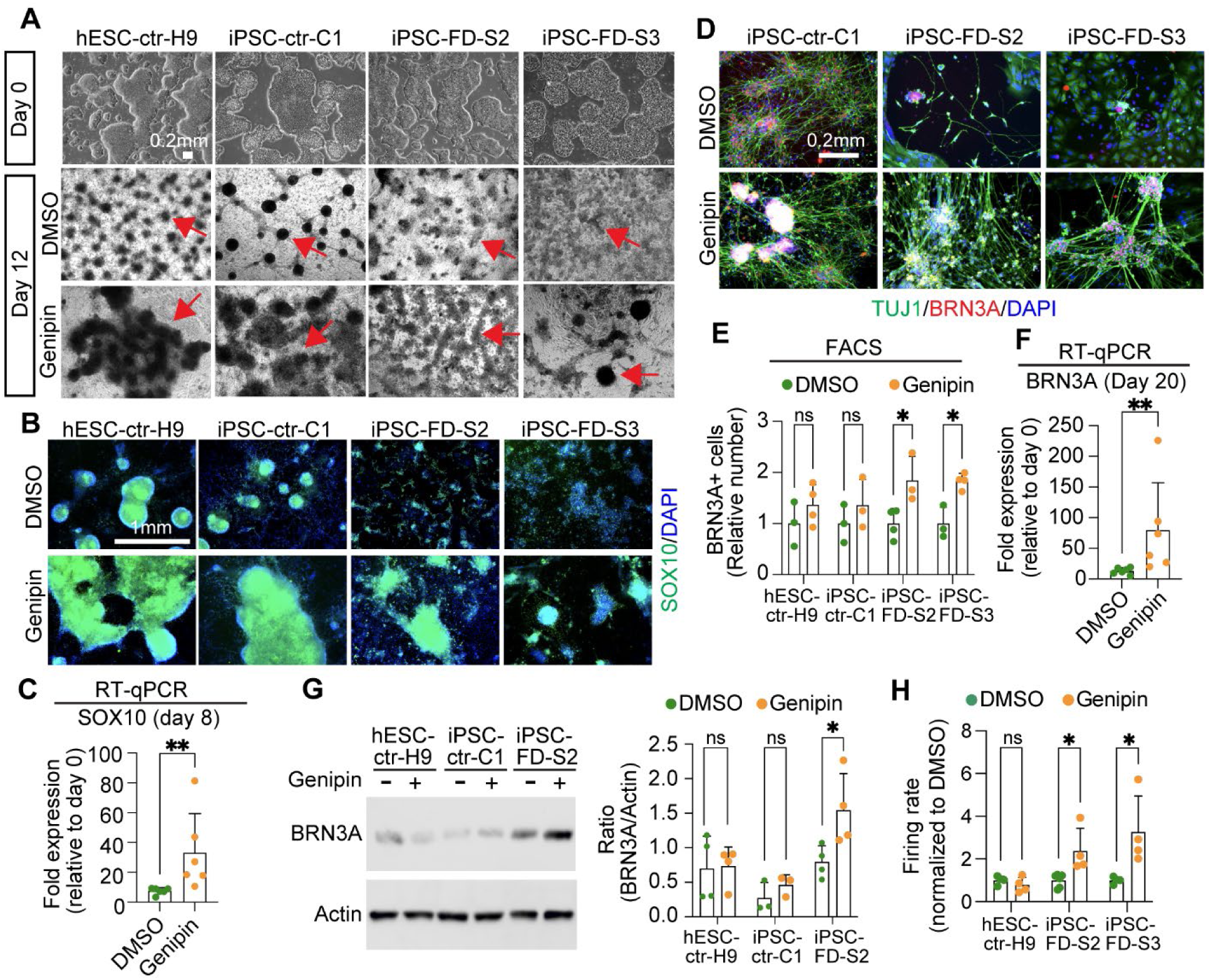
Genipin rescues neural crest and sensory neuron-related phenotypes in FD. **A)** hPSC lines look normal at the pluripotent stage (day 0), but differentiation into NC cells is diminished in the FD lines (DMSO, dark ridges/red arrow indicate NC cells); this is rescued by genipin (10 µM) treatment. **B and C)** SOX10 expression is restored in NC cells upon genipin treatment. Cells were differentiated in the presence of genipin (10 µM) and fixed on day 12. **B)** Cells were stained for SOX10 and DAPI and analyzed by IF. n=5 biological replicates. **C)** Genipin increases SOX10 expression. RNA isolated from FD cells differentiated in the presence of genipin (10 µM) was analyzed by RT-qPCR. n=6 biological replicates. **D-F)** Genipin restores SN differentiation. iPSC-FD-S2 and iPSC-FD-S3 cells were treated with genipin (10 µM) and differentiated into SNs. RNA was isolated on day 20 or cells were fixed and stained using the indicated antibodies and analyzed by **D)** IF (n=5 biological replicates), **E)** intracellular flow cytometry (n=3-4 biological replicates), and **F)** RT-qPCR (n=6 biological replicates). **G)** Western blot analysis confirms the increase in SN production upon genipin treatment. Cells differentiated with genipin (10 µM) were lysed on day 20 and immunoblotted with the indicated antibodies (left) and quantified (right). n=3-4 biological replicates. **H)** Genipin increases firing rate of FD SNs. Cells were differentiated in the presence of genipin (10 µM) and firing rate was analyzed by MEA. n=4-6 biological replicates. In **C, E, F, G, H**, Two-tailed Student’s t-test. ns, non-significant, *p<0.05, **p<0.005. All graphs show mean ± s.d. Data from iPSC-FD-S2 and iPSC-FD-S3 are pooled as FD in **C** and **F**.

### Genipin rescues sensory deficits in FD mice

We next assessed whether genipin can also rescue the FD neurodevelopmental defects *in vivo*. Since FD mutant mouse embryos (*Elp1*^Δ20/flox^) have significant deficits already at late gestation^9^, we assessed whether genipin has the potential to rescue their developmental defects. Pregnant dams were treated with genipin-supplemented chow from mid-gestation. At E18.5 embryos were harvested and analyzed (**Fig. 4A**). Genipin was well tolerated during gestation and no significant effect was observed in female gestational weight gain, litter size, or embryonic development (**Sup. Fig. 2F** and **Sup. Table 3**).

**Figure 4.**
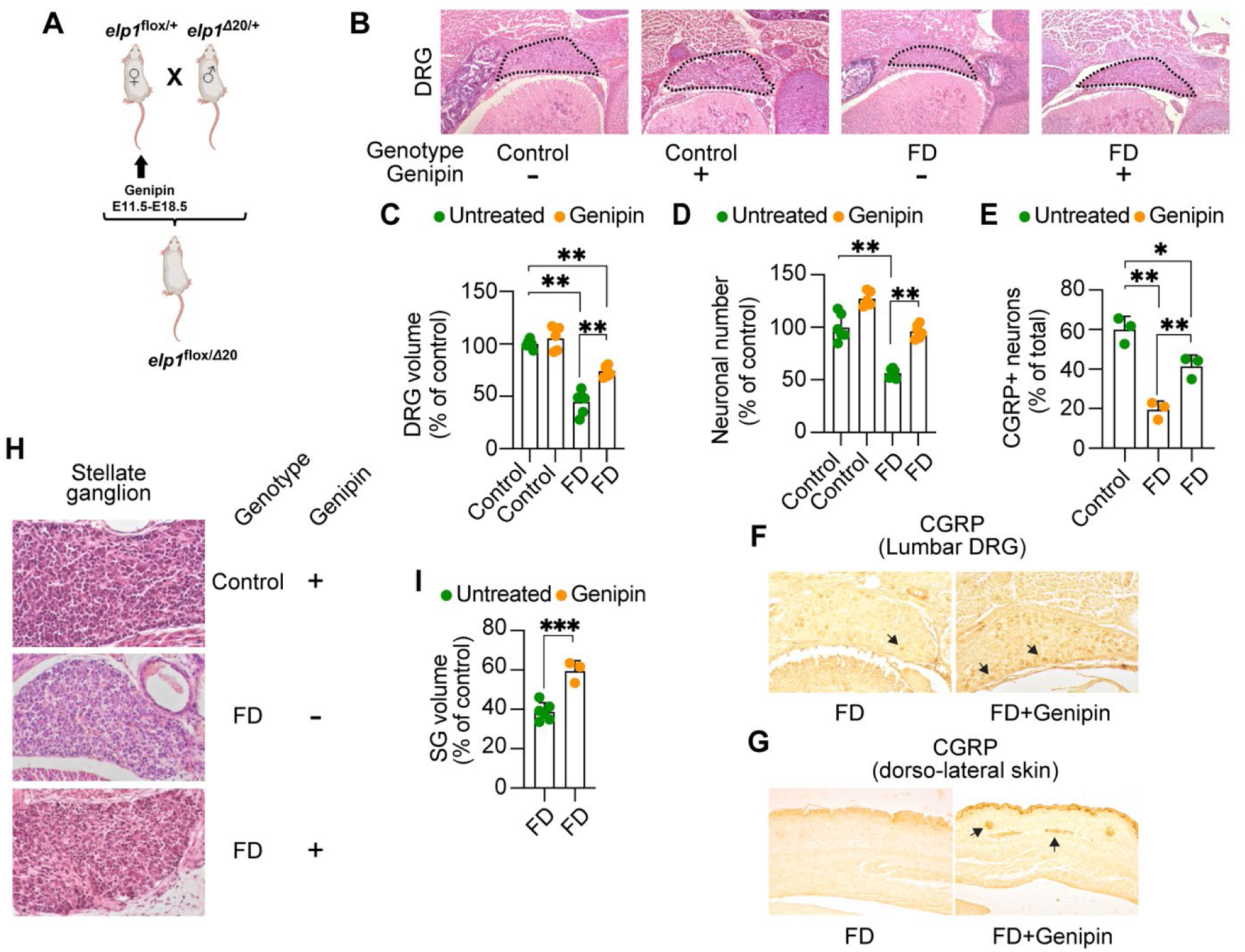
Genipin rescues FD peripheral deficits *in vivo*. **A)** Breeding and treatment schematic. **B)** Representative H&E-stained transverse sections through lumbar (L1) dorsal root ganglia (DRG) of untreated control, genipin-treated control, untreated FD, and genipin-treated FD E18.5 embryos, at their largest dimensions. **C)** Volumes of lumbar (L1) DRGs of untreated (n=6) and genipin-treated (n=5) controls, and untreated (n=6) and genipin-treated (n=6) FD E18.5 embryos, displayed as percentage of control. **D)** Neuronal counts of lumbar (L1) DRGs of untreated (n=6) and genipin-treated (n=5) controls, and untreated (n=6) and genipin-treated (n=6) FD E18.5 embryos, displayed as percentage of control. **E)** Percentage of CGRP-positive neurons in lumbar (L1) DRGs of control untreated (n=3), FD untreated (n=3), and genipin-treated FD (n=3). **F)** Representative images of transverse sections through lumbar (L1) DRGs of untreated FD and genipin-treated FD E18.5 embryos immunostained with CGRP. **G)** Representative images of transverse sections through dorso-lateral skin of untreated FD and genipin-treated FD E18.5 embryos immunostained with CGRP. Arrows show positive signal. **H)** Representative H&E-stained sections of stellate ganglia (SG) of genipin-treated control, untreated FD, and genipin-treated FD E18.5 embryos. **I)** Volumes of SG of untreated (n=6) and genipin-treated (n=3) FD E18.5 embryos, plotted as percentage of control. All graphs show mean ± s.d.. For **C, D, E**, one-way ANOVA followed by Tuckey HSD post hoc test. For **I**, two-tailed t-test. *p<0.05, **p<0.01, ***p<0.001

At E18.5, FD embryos treated with genipin did not differ significantly in weight and size compared to untreated FD embryos (779.33 mg ± 94.00 versus 783.6 mg ± 137.62, respectively) but remained significantly smaller than control embryos^9^. On the other hand, genipin treatment during gestation significantly rescued FD DRG volume and neuronal cell numbers (**Fig. 4B-D**). Since sensory nociceptive neurons are particularly depleted in FD at late gestation^9, 29, 30^, we assessed the paucity of nociceptive neurons by immunohistochemistry for calcitonin gene related protein (CGRP), a specific marker for nociceptive neurons^31^. As shown in **Fig. 4E** and **F**, while in FD embryos CGRP-positive neurons are reduced to 20% of total lumbar L1 DRG neurons, in genipin-treated FD embryos the number of CGRP-positive neurons increased significantly to about 40% of total neurons. Interestingly, we found that genipin treatment also rescued sensory skin innervation in FD mouse embryos (**Fig. 4G**), which is severely compromised in FD patients and FD mouse models^29, 32^. Unexpectedly, we found that genipin also rescued neurodevelopmental phenotypes in sympathetic neurons, assessed by an increase in the size of the stellate ganglion (**Fig. 4H, I**).

These results suggest that genipin is well tolerated and rescues FD sensory and sympathetic phenotypes *in vivo*. Furthermore, they confirm the results we obtained in the hPSC-based system and strengthen our hypothesis that genipin is a suitable candidate for as a therapeutic to treat neurodevelopmental diseases such as FD.

### Effects of genipin on neurodegeneration in sensory neurons in FD

We have previously shown that SNs in FD degenerate and die over time *in vitro* in our PSC model^17^. This correlates with reports in FD patients of deteriorating symptoms with age, including pain perception and autonomic regulation^18^. Since FD is both a neurodevelopmental and a neurodegenerative disorder and thus patients are born with reduced numbers of SNs, it would be imperative for patient care to develop a novel drug that halts or slows degeneration of existing SNs. Therefore, we set out to assess genipin’s capability to halt SN degeneration. We first had to adapt our *in vitro* degeneration assay^17^ to the new SN differentiation protocol^13^. Titration of NGF revealed that 1 ng/ml of NGF leads to degeneration of iPSC-FD-S2 SNs over 20 days while being able to maintain healthy SNs alive (**Sup. Fig. 3A**, red rectangles). Also, exclusion experiments showed that LM, but not FN or PO could be omitted in the surface coating for SNs (**Sup. Fig. 3B**). Thus, survival assays were done in 1 ng/ml NGF on dishes coated with PO and FN. SNs were differentiated from hPSC lines without genipin until day 12, when they were replated and treatment with genipin began (**Fig. 5A**). We found that, healthy control SNs survive well *in vitro* until day 34 and beyond, and they show the classical clustering/ganglia-like structure formation over time (**Fig. 5B**, arrows). Severe FD SNs, however, die starting from day 13 (**Fig. 5B, C**). When the SNs are treated with genipin from day 13, this degeneration is prevented (**Fig. 5B, C**). We confirmed this effect in the KSR-based protocol (**Fig. 1A, Sup. Fig. 3C**). Furthermore, we show that neurite length from FD SNs increases compared to untreated SNs (**Fig. 5D, E**). We also found that genipin increased the number of dendrites of FD SNs and renders a distribution more similar to healthy SNs (**Sup. Fig. 3D**). It is of note that in a 2D *in vitro* system SNs show more neurites compared to *in vivo*^33^. To make sure that there are not substantial numbers of NC cells remaining at day 13, that could be induced by genipin to make more SNs, we stained day 13 and day 34 cultures for *SOX10*, showing the largely absence of NC cells (**Sup. Fig. 3E**). Thus, genipin truly supports survival of SNs in FD. Together, these results suggest that genipin is a promising candidate drug to halt the progressive SNs degeneration in FD and further might become an interesting candidate drug for the treatment of other peripheral neuropathies.

**Figure 5.**
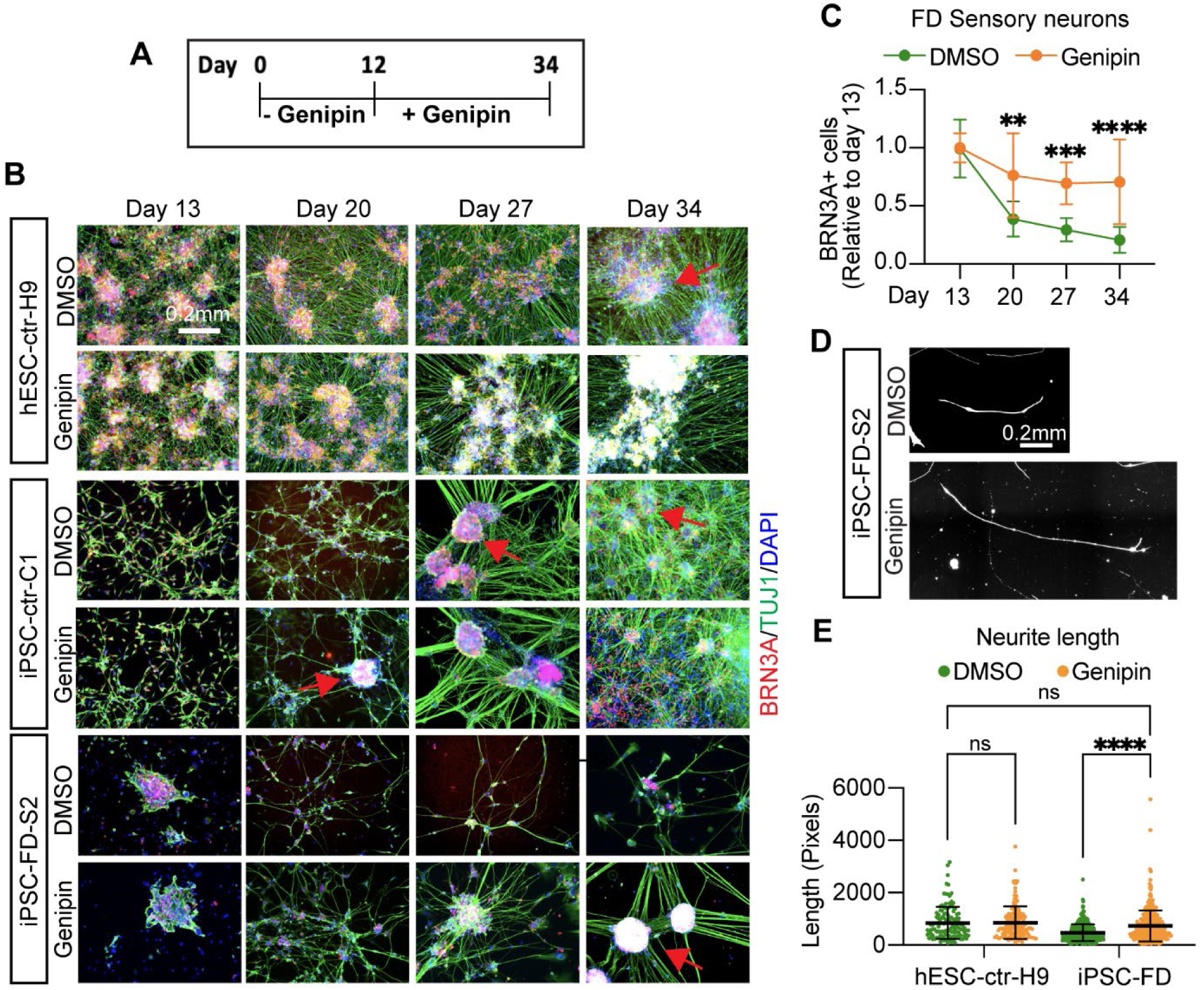
Genipin rescues sensory neuron degeneration in FD. **A)** Schematic of survival assay. **B)** Healthy or FD cells were treated with genipin from day 12 on (when neurons are born) and monitored for survival for 21 days. Cells were fixed on the indicated days and stained for BRN3A, TUJ1 and DAPI. Arrows indicate healthy, ganglia-like SN clusters. **C)** Image quantification of **B**. n=8, Two-way ANOVA followed by Šídák multiple comparisons. **p<0.005, ***p<0.001, ****p<0.0001. **D,E**, Genipin increases neurite length and number in FD cells. **D)** Representative images of neurite length. **E)** Measurement of neurite length (number of pixels) of **D**. n=6-8 biological replicates, one-way ANOVA followed by Tukey’s multiple comparisons. ns, non-significant, ****p<0.0001. All graphs show mean ± s.d. Data from iPSC-FD-S2 and iPSC-FD-S3 are pooled as FD in **C**, and **E**.

### ELP1 targets ECM gene expression

We next addressed the mode of action through which genipin exerts these rescue effects. FD is caused by a homozygous mutation in *ELP1*, leading to a splicing defect and eventually causing patients to have reduced levels of wild type ELP1 protein, with particularly low levels in PNS tissues, including SNs^6^. Several reports have shown compounds that reverse this splicing defect^34–37^, thus we first assessed if genipin directly affects ELP1. We found that genipin does not change splicing of *ELP1*, assessed on the mRNA and on the protein level (**Sup. Fig. 4A, B).** Thus, in FD at the SN level genipin must exert its action in a different way.

Genipin has been implicated in several contexts, including inhibition of mitochondrial protein UCP2, activation of neuronal nitric oxide synthetase (NOS)^20^ and its ability to crosslink ECM proteins^38^. Our previous work ^17^ has shown that severe FD patients have a modifier mutation in *LAMB4*, which is an ECM protein. Furthermore, proper composition and availability of ECM proteins have been shown to be essential for SN development^39–41^. Additionally, Goffena et al. showed that loss of Elp1 particularly affects long transcripts and AA- and AG-ending transcripts^42^. We analyzed their sequencing data and found that ECM proteins fall within this highly affected group (**Sup. Fig. 4C, D**). Furthermore, we conducted RNA sequencing analysis comparing SNs from healthy control and iPSC-FD-S2 cells. Gene ontology analysis revealed that ECM proteins significantly downregulated in FD SNs (**Sup. Fig. 4E**). Further analysis of gene differential expression showed that the ECM-related proteins *FBLN1*, *COL1A2*, *HAPLN1*, *TNC*, and *HSPG2*, among others are downregulated (**Sup. Fig. 4F**). Thus, loss of *ELP1* is predicted to lead to ECM defects and we hypothesized that genipin’s ability to crosslink ECM proteins might alleviate these issues in FD.

### Genipin acts via crosslinking of extracellular matrix proteins in Familial Dysautonomia

Genipin forms inter- and intramolecular crosslinks (**Fig. 6A**), which cause the cells to be stained blue due to formation of genipin-methylamine monomers during crosslinking reactions (**Fig. 6B**). Thus, we reasoned that if the rescue in FD functions via ECM crosslinking, then other crosslinking agents should have the same rescue effect as genipin in our FD model. We first tried dimethyl pimelimidate (DMP), a membrane-permeable compound that crosslinks crosslink primary amines of proteins (due to imidoester groups present on each end of the molecule) both within the ECM and inside the cell (**Sup. Fig. 5A**). We found that DMP rescued both the NC deficit (**Sup. Fig. 5B**, top) as well as the SN deficit (**Sup. Fig. 5B**, bottom) in FD cells in a dose dependent manner (**Sup. Fig. 5C**). This result suggests that protein crosslinking is sufficient to rescue the neurodevelopmental FD phenotype. However, it remains unclear whether crosslinking of extracellular or intracellular proteins or both rescue the FD phenotype. Thus, we used bis-(sulfosuccinimidyl) suberate (BS3), a membrane-impermeable crosslinker which only crosslinks proteins in the ECM (**Fig. 6C**). We found that BS3 rescues both the NC (**Fig. 6D**, top) as well as the SN phenotypes (**Fig. 6D**, bottom) in a dose dependent manner (**Sup. Fig. 5D**). Additionally, when we used N-hydroxysulfosuccinimide (Sulfo-NHS), a form of BS3 with no crosslinking capabilities, the rescue effect was abolished in both the NC and SN phenotypes (**Sup. Fig. 5E**). Together, our data suggests that crosslinking of only extracellular proteins (i.e. the ECM) is necessary and sufficient to rescue the developmental phenotype in FD. Moreover, these results strongly support a model where genipin exerts its action in FD via crosslinking proteins in the ECM. To confirm that the FD phenotype is rescued only by genipin’s crosslinking activity and not its other predicted function (i.e. UCP2 inhibition), we used 1,10-anydrogenipin (AG, **Fig. 6E**). AG has been shown to specifically remove genipin’s ability to crosslink proteins, while keeping its other actions intact^24^. Confirming that AG does not have crosslink activity, we found that AG did not stain the FD cells blue (**Fig. 6F**). Furthermore, in contrast to genipin, AG was not able to rescue the NC (**Sup. Fig. 5F, G**) nor the SN phenotype in FD cells (**Fig. 6G,H, Sup. Fig. 5H, I**). Together, our results show that genipin’s crosslinking action is responsible for the FD phenotype rescue.

**Figure 6.**
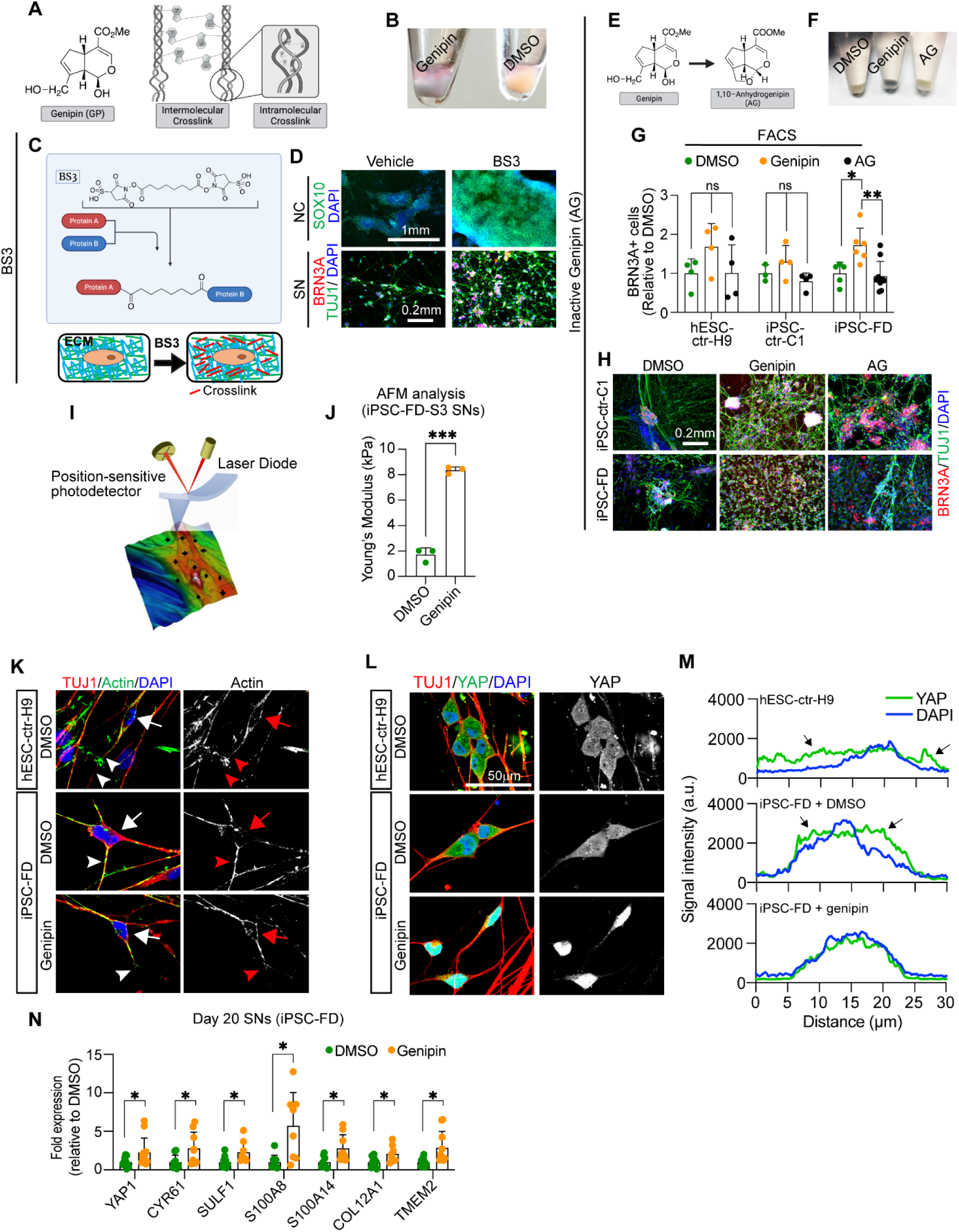
Genipin rescues FD phenotypes via crosslinking of extracellular matrix proteins and activates. **A)** Genipin chemical structure and schematic of intermolecular and intramolecular crosslinking. **B)** Treatment with genipin turns cells blue. **C)** Schematic of BS3 crosslinking action and its extracellular location. **D)** BS3 rescues the NC and SN differentiation defect in FD. FD cells were differentiated in the presence of DMP and fixed on day 12 (NC) and day 20 (SN). Following staining using the indicated antibodies. n=3-5 biological replicates. **E-H)** 1,10-anhydrogenipin (AG) does not rescue the SN defect in FD. **E)** AG chemical structure. **F)** AG does not turn cells blue. **G, H)** AG does not promote FD SN differentiation. Cells were differentiated into SNs in the presence of genipin or AG. SNs were fixed on day 20 and stained for BRN3A for intracellular flow cytometry analysis (**G**, n=6-8 biological replicates), or stained for BRN3A, TUJ1, and DAPI for IF (**H**, n=6-8 biological replicates). **I, J)** Genipin-mediated crosslinking increases ECM stiffness. **I)** AFM Experiment schematics. **J)** Genipin increases the Young’s modulus of SNs. iPSC-FD-S3 SNs were fixed on day 25 and analyzed by AFM (n=3 biological replicates). **K-M)** Genipin reorganizes the actin cytoskeleton and induces transcription of YAP-dependent genes. **K)** Genipin partially rescues the differences of actin expression pattern in healthy and FD SNs. Day 20 SNs were fixed and stained for the indicated antibodies. Images were obtained by confocal microscopy. Actin signal in the cell body (arrows) or the axons (arrowheads) are highlighted. **L, M)** YAP localization changes in the presence of genipin. Day 20 SNs were fixed and stained for the indicated antibodies. **M)** YAP and DAPI signal intensity from images on **L)** was measured and plotted n=6-7 biological replicates). Arrows indicate YAP signal outside of the nucleus (stained with DAPI). **N)** Expression of YAP-dependent genes. RNA from FD SNs treated with DMSO or Genipin was extracted on day 20 and gene expression was analyzed by RT-qPCR (n=7-9 biological replicates). All graphs show mean ± s.d.. For **G**, one-way ANOVA followed by Tukey’s multiple comparisons. For **J**, two-tailed t test with Welch’s correction. For **N**, two-tailed t test. ns, non-significant, *p<0.05, **p<0.005, ***p<0.001. In **G, M,** and **L** data from iPSC-FD-S2 and iPSC-FD-S3 are pooled as FD.

### Genipin increases stiffens of the ECM of FD SNs

Atomic force microscopy (AFM) can be used to study how the morphology and mechanical properties of live or fixed cells are affected under different conditions (for example, changes in the ECM)^43–45^. In AFM, a tip is roughly aligned on the surface of the cell using an optical microscope, then contact AFM mode is applied where the tip scans the surface (**Fig. 6I**). A laser beam, aimed at the cantilever holding the tip, is detected by a photodetector. Vertical changes in the position of the tip (due to different height in the surface) are measured, resulting in high-resolution topographic images of the samples (**Fig. 6I, Sup. Fig. 6A**). Interestingly, we found that genipin re-orientates the ECM as shown by the presence of ordered “strips” (possibly due to the presence of crosslinking bundles), in stark contrast to DMSO controls where the ordered “strips” are missing (**Sup. Fig 6A**). Also, the height of the samples increased with genipin (from to 2.09 µm to 3.5 µm for SNs, from 0.8 µm to 1.2 µm for ECM) (**Sup. Fig 6A**) due to formation of crosslinked bundles. From these acquired cell images, we selected different points on the surface (**Fig. 6I**, black crosses) and measured their force spectroscopy. The deflection of the cantilever tip was recorded and converted to force-distance curves (**Sup. Fig. 6B**, solid lines). Force-distance curves were further fitted with the Hertz model (suitable for fitting under small deformation in nanoindentation measurements^46^) to calculate the Young’s modulus (a measurement of stiffness) of individual cells (**Sup. Fig. 6B**, dotted lines). We found that FD SNs treated with genipin have a higher Young’s modulus (∼8,000 Pa) compared to DMSO control (∼2,000 Pa) (**Fig. 6J, Sup. Fig 6B, C**), agreeing with previous reports^45, 47, 48^. Furthermore, genipin-treated SNs show narrower distributions, suggesting a stiffer and more uniform surface due to ECM crosslinking (**Sup. Fig 6D**). To confirm our results, we removed the SNs and measured only the remaining ECM. Agreeing with our previous results, we found that the ECM deposited by genipin-treated SNs has a Young’s Modulus ten times higher compared to untreated controls (∼13,300 Pa vs ∼1,300 Pa) (**Fig. 6J, Sup. Fig 6B, C**). These changes could only be attributed to the direct crosslinking of genipin with ECM, further strengthening our hypothesis that ECM crosslinking by genipin is necessary to rescue the FD neurodevelopmental and neurodegenerative phenotypes.

### ECM crosslinking causes actin reorganization and transcription of YAP-dependent genes

Based on our previous results, we asked whether a stiffer ECM (due to genipin-dependent crosslinking) has any intracellular effects. The ECM is necessary for proper development and distribution of the cytoskeleton^49^, and previous reports showed that the actin cytoskeleton is missregulated in FD^50^. Thus, we first assessed whether the actin expression pattern is different between healthy and FD SNs. We measured the signal from the cell bodies (**Sup. Fig. 6E**, black line) or the axons (**Sup. Fig. 6E**, blue lines). We found that the actin signal in healthy SNs is the highest in the cell body, whereas FD SNs show stronger actin signal in the axon (**Fig. 6K, Sup. Fig. 6F**, **G**). Treatment with genipin was able to partially restore the actin expression pattern in FD SNs (**Fig. 6K**). Changes in actin cytoskeleton organization due to ECM stiffness activate the Hippo pathway^51^. In a stiff ECM, the transcriptional coactivator YAP (the most downstream effector of the Hippo pathway), translocates from the cytoplasm to the nucleus to activate a transcriptional program^51^. We sought to assess YAP localization in SNs by immunofluorescence. We found that in both healthy and FD SNs, YAP is present in the cytoplasm as assessed by the presence of YAP signal outside of the nucleus (stained with DAPI) (**Fig. 6L, M**, black arrows). However, upon genipin treatment, YAP is transported to the nucleus as shown by detection of YAP fluorescent signal only within the boundaries of the nucleus (stained with DAPI) (**Fig. 6L, M**). This suggests that the increase in ECM stiffness due to genipin-mediated crosslinking activates the Hippo pathway in SNs. We next assessed whether YAP nuclear translocation results in transcription of YAP-dependent genes. Indeed, we found that *YAP1* and *CYR61* expression is significantly higher in FD SNs treated with genipin (**Fig. 6N**). RUNX1 is a transcription factor necessary for maturation of nociceptors (a subtype of SNs). Interestingly, in cancer YAP interacts with RUNX1 to regulate transcription^52, 53^, thus we asked whether these genes are also activated in FD SNs in the presence of genipin. We found that genipin increased the expression of all the tested genes that are dependent on YAP-RUNX1 interaction: *SULF1*, *S100A8*, *S100A14*, *COL12A1*, and *TMEM2* (**Fig. 6M**). Together, our data suggest that genipin-dependent ECM crosslinking causes actin reorganization and activation of the Hippo pathway in SNs, which rescue development and survival in FD phenotypes.

### Genipin enhances axon regeneration of neurons from the peripheral and central nervous system

Our results show that genipin rescues both developmental and degenerative FD phenotypes in SNs, suggesting its beneficial support of neurons in this disease. We wondered whether genipin might be beneficial in more common disorders of the PNS. To test this hypothesis, we inquired if genipin is beneficial for axon regeneration. We first assessed whether we can use our *in vitro* system as an axotomy model. We have previously shown that in our system, hPSC-ctr-H9-derived SNs can be replated at late stage^13^, which is known to enzymatically cut neurites and thus serving as an axotomy model (**Fig. 7A**). Employing this approach, in combination with genipin treatment during the regeneration phase, we found that this process causes expression of *ATF3* and *SPRR1A*, two critical genes involved in neuron regeneration^54^, in DMSO treated SNs differentiated from hPSC-ctr-H9 cells (**Fig. 7B**). Interestingly, we found that SNs replated in the presence of genipin express higher levels of both genes (**Fig. 7B**). Furthermore, high *ATF3* expression is maintained for a longer period of time. We next assessed whether genipin affects the speed of axon regeneration. We found that SNs treated with genipin showed longer axons 18 hours and 6 days post replating (**Fig. 7C, D**). To expand genipin’s application in the PNS further, we tested peripheral sympathetic neuron regeneration. To do so, we employed a microfluidic device, which maintains cell bodies and axons in two different compartments, allowing us to break the axons without disturbing the cell bodies (**Fig. 7E**). An NGF gradient guides the direction of axonal growth and neurites can be cut and removed with a pipet tip from the left chamber without hurting the cell bodies in the right chamber (**Fig. 7E**). We found that genipin increases the number and length of regenerated axons from hPSC-ctr-H9-derived sympathetic neurons 2 days post injury, measured by an increase in PRPH staining by immunofluorescence (**Fig. 7F, G**). These results suggest that our system can be used to model axonal injury in the PNS and that genipin increases the regeneration speed in peripheral neurons. Finally, we asked if genipin might be beneficial for neurite regeneration in CNS neurons, possibly extending its usefulness further to spinal cord injury in a broader context. To do that, we tested prefrontal cortical (PFC) neurons differentiated from hPSC-ctr-H9 cells^55^. Using our replating method (**Fig. 7H**), we found that PFC neurons treated with genipin regenerate their axons faster than DMSO-treated neurons by IF, measured by the increase in the number of pixels positive for TUJ1 (**Fig. 7I-J**). Taken together, our results suggest that genipin can be used to promote neuronal regeneration in both PNS and CNS neurons applicable to a broad range of degenerative diseases.

**Figure 7.**
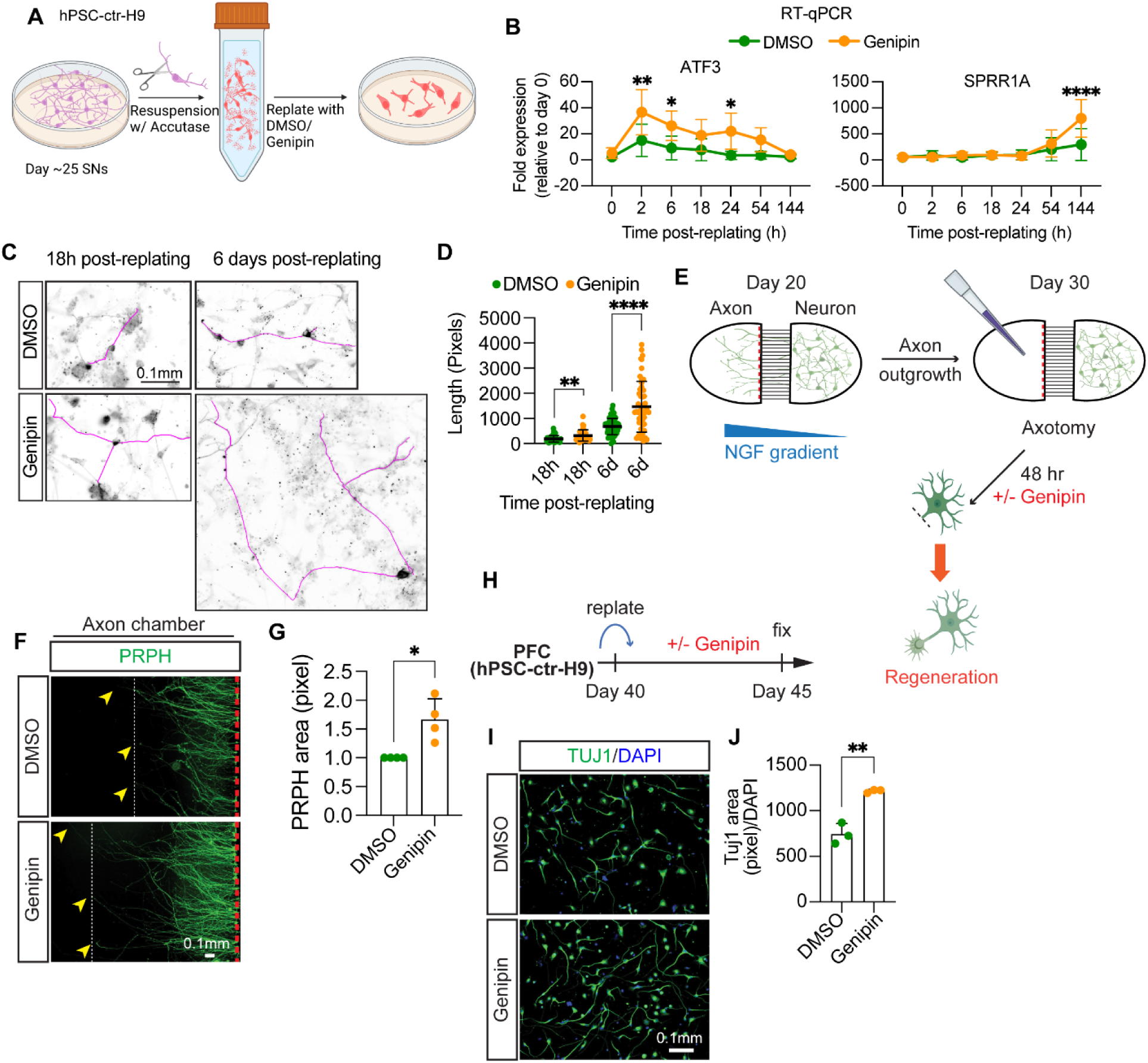
Genipin accelerates axon regeneration after injury in different neuronal types. **A-D)** Genipin enhances rengeneration of healthy SNs. **A)** Experiment schematics. **B)** Genipin increases expresison of injury-related genes. RNA from hPSC-ctr-H9 SNs was isolated at indicated times and gene expression was measured by RT-qPCR (n=4-7 biological replicates). **C**,**D)** Genipin increases axon length after injury. Day 25 SNs from hPSC-ctr-H9 cells were fixed at indicated times after replating in the presence of DMSO or Genipin, and stained for TUJ1. Axons were traced (magenta) and measurement were plotted in **D** (n>20 cells from 5 biological replicates). **E-G)** Genipin increases length and complexity of axons from sympathetic neurons. **E)** Experiment schematics. For details, see Methods. **F,G)** Axons from sympathetic neurons were removed. After 48 h treatment with Genipin, neurons were fixed and stained for PRPH. Axons were measured (arrowheads indicate length of axons) and the pixels with high PRPH signal were measuered and graphed in **J** (n=4 biological replicates). **H-J)** Genipin promotes axon regeneration in prefrontal cortical (PFC) neurons. **H)** Experiment schematics. For details, see Methods. **I,J)** hPSC-ctr-H9 PFC neurons reaplted and incuabted with Genipin for 5 days. Neurons were then fixed and stained for TUJ1 as shown in **I**. Measurement of the pixels with high TUJ1 signal were normalized to DAPI and plotted in **J** (n=3 biological replicates). All graphs show mean ± s.d. For **B**, two-way ANOVA followed by Šídák multiple comparisons. For **D, G**, and **H**, two-tailed t test. *p<0.05, **p<0.005, ***p<0.001, ****p<0.0001.

## Discussion

Here, we discovered that genipin rescues both neurodevelopmental and neurodegenerative phenotypes in a hPSC-based disease model of FD. We further showed that genipin rescues sensory ganglia phenotypes *in vivo*. Additionally, we demonstrate that these phenotypes are due to crosslinking of ECM proteins causing rearrangement of the cytoskeleton and activates transcription of YAP-dependent genes. Finally, we showed that genipin promotes regeneration of neurons from the peripheral and central nervous system after injury. Thus, genipin has a broad application potential in a wide range of neurological disorders.

To date there is a lack of treatments that specifically target either the neurodevelopmental or the neurodegenerative aspects of FD. This void leads to patient’s symptoms being managed both surgically and pharmacologically^2, 10^ without clearly targeting the molecular cause of the disease. Over the past 20 years, multiple compounds have been identified that target the ELP1 splicing defect in FD have been identified, however they have failed in clinical trials or there is no evidence that they target the PNS. These include the plant hormone kinetin, which, despite its promising results in FD animals^34^, worsened the nausea and vomiting episodes in patients^2^. Phosphatydilserine (PS) which increased ELP1 expression in patient-derived fibroblasts^56^ and animal models^57^ but has not successfully undergone clinical trials^2^. Finally, tocotrienol (vitamin E) and the antioxidant epigallocatechin gallate (ECGC) were increase *ELP1* expression in cells lines^58,59^but did not increase *ELP1* expression in patients, nor improved clinical outcomes^16, 60, 61^. A recent study showed that the small molecule RECTAS restores normal *ELP1* splicing in SNs differentiated from FD iPSCs and *in vivo*, however it is unclear whether it rescues any FD phenotype^37^. Finally, gene therapy approaches, such as antisense oligonucleotides (ASO), have been shown to restore ELP1 expression in patient fibroblasts and FD animals^62–64^. Importantly, however, they did not show a rescue in PNS tissues and human clinical trials are note initiated yet. In sum, the field lacks specific drugs to treat FD and the available candidates have not been tested or are not effective in human tissues affected in patients, such as PNS neurons.

Similar to FD, more common neurodegenerative disorders affecting the PNS experience a dramatic absence of available treatments. These include diabetes (in up to 50% of patients) or chemotherapy-induced (in up to 40% of patients) peripheral neuropathy, as well as neuropathies caused by injury, infections, medications, alcoholism, and even vitamin deficiencies. In fact, there are very few FDA-approved drugs available for peripheral neuropathy and PNS injury and the ones available mainly focus on managing symptoms^11, 65^. Furthermore, although there is extensive research aimed to identify treatment that regenerate peripheral neurons after injury, very few of them are or have been in clinical trials. They include gene therapy, electrical stimulation, and nerve transfer of an expendable, uninjured donor nerve^66^. Due to these difficulties, genipin could be a cost-effective treatment for nerve regeneration. Our results show that genipin is a highly promising compound with potential to be further developed as a treatment option for neurodevelopmental and neurodegenerative diseases (including peripheral neuropathies). Additionally, its lack of toxicity on cells^26^ and *in vivo*^24, 25^, and the fact that it can transverse through the placenta, makes genipin a safe drug. Further studies will be needed to address this.

Our results showing genipin’s ability to rescue neurodevelopmental defects in hPSC-derived FD SNs and in SNs in mice highlight the importance of ECM proteins during development. This data lays the groundwork to further investigate the ECM composition necessary for normal PNS development, as well as how defects in ECM proteins, such as laminin^17^, contribute to PNS disorders^67^. The rescue on neurodevelopment that we see in FD mice is interesting and urges the investigation of the effects of genipin on neurodegeneration further. As we report, genipin reverses the progressive loss of SNs in the FD hPSC-based disease model and it enhances neurite growth and density (**Fig. 5B-E, Sup. Fig. 3D**). This is consistent with the literature, where it has been shown that the extracellular environment (presence of growth factors, ECM composition, and 2D vs 3D cultures) affect neurite length and arborization^33, 40, 68, 69^. This is in agreement with the role of genipin as an ECM crosslinker. It will be crucial to assess this finding further, as this may provide real promise for patients with neurodegenerative diseases. For instance, in chemotherapy-derived neuropathies, patients who are more susceptible to axonal degeneration are affected by paclitaxel-induced neuropathy^74^. Genipin could be used as a supplement to prevent neuronal degeneration.

Most research on the molecular implications of FD focus primarily on *ELP1*. We show that genipin exerts it effects via crosslinking of ECM proteins and that its rescue effect is removed upon removal of its crosslinking activity (**Fig. 6E-H, Sup. Fig. 5F-I**). Agreeing with this, we show that genipin increases surface stiffness of SNs and isolated, deposited ECM proteins (**Fig. 6J, Sup. Fig. 6B-D**). Previous reports, showed that loss of ELP1 results in defective actin and microtubule organization in fibroblasts or brain tissues^50, 70^. Here, we report that these phenotypes are conserved in SNs differentiated from human FD iPSCs (**Fig. 6K, Sup. Fig. 6F, G**). Furthermore, we found that increasing stiffness of the ECM by crosslinking rescues this phenotype, agreeing with studies showing that biomechanical cues (such as stiffness of the ECM) affect actin cytoskeleton organization^71^. Our RNAseq results show that upon loss of *ELP1* a myriad of ECM-related proteins are downregulated causing an aberrant cellular environment. Therefore, signaling multiple pathways are missregulated. One of them is the Hippo pathway, which translates external mechanical cues into biochemical signals that affect gene expression^51^. Interestingly, we show that YAP (the most downstream effector of the Hippo pathway) is mainly present in the cytoplasm of SNs, suggesting that it is inactive, agreeing with previous reports^72^. However, upon genipin treatment, YAP translocates to the nucleus and promotes transcription of genes associated with cell proliferation (*CYR61*, *S100A14*), cell cycle progression (*S100A8*), ECM remodeling (*SULF1*, *COL12A2*, *TMEM2*) (**Fig. 6L-N**). We hypothesized that YAP activation initiates a transcriptional program that allows the proliferation of progenitor cells, thus increasing the number and survival of SNs. Very little is known of the role of the Hippo pathway and YAP function in the PNS, although mechanotransduction (including ECM-cell interaction) is important in NC migration and differentiation^67^. Furthermore, it has been proposed that mechanical modulation of the Hippo pathway could potentially be used as a treatment for peripheral neuropathies^73^. Genipin, by its ECM-crosslinking activity, fit in this category. Our results show that potential treatment options for FD lay beyond ELP1 splicing correction and that modifying the ECM may be key. Furthermore, our findings suggest that looking at basic cellular mechanisms is necessary to identify new potential therapeutic targets.

In sum, genipin shows great promise as a future treatment option for FD as well as other more common neuropathies. Its action via ECM crosslinking and our exciting results about its ability to increase axon regeneration make it an interesting compound for the future application to nerve regeneration in the PNS and possibly prevention of peripheral neuropathies.

## Materials and methods

### Chemical screen

For the chemical screen hPSC-ctr-H9 control and iPSC-FD-S2 cells were cultured in KSR conditions (see below). 96-well plates were coated with Matrigel. hPSCs were detached from MEF cultures using Trypsin (52) and cells were seeded at 80,000 cells/well for iPSC-FD-S2 and 40,000 cells/well for hPSC-ctr-H9 in order to start the differentiation at equal cell numbers the following day (=day 0). The SN differentiation was performed as described in Fig. 1c. The compounds of the LOPAC chemical library (Sigma) were added on differentiation days 2, 5 and 8 and the plates were fixed and stained for BRN3A and DAPI on day 10. 16 tiles/well were imaged using the MetaXPress software: Cell Scoring by Molecular Devices. The percentage of BRN3A+ cells over DAPI of all DMSO control wells was averaged. Each compound-treated well number was divided by this DMSO average to calculate the fold change/fold increase of compound-treated wells over controls. To call a hit the average + 3 SD was used. The Z’ score (53) (Z’=0.25) was calculated before conducting the screen, using a pilot screen with only hPSC-ctr-H9 and iPSC-FD-S2 lines in a small format, but in the final screening conditions. The Z-score for genipin at 1 mM was 15 and at 10mM it was 17.

### hPSC maintenance

In KSR conditions, hPSCs were maintained on MEFs as described^17^. In E8 conditions, human embryonic stem cells (WA-09, WiCell) and all iPSC lines were grown on dishes coated with vitronectin (5μg/mL, 1h at RT) at 37°C with 5% CO2 and fed daily with Essential 8 Medium + Supplement. For splitting, hPSC-ctr-H9 colonies were washed with PBS and incubated with 0.5mM EDTA, 3.08M NaCl in PBS with for 2 min at 37°C. Cells were then resuspended in E8 + Supplement and split at a ratio of 1:10. iPSC-ctr-C1, iPSC-FD-M2, iPSC-FD-S2, and iPSC-FD-S3 cells were previously characterized ^17^ and maintained under the same conditions.

### Neural crest and sensory neuron differentiation

In KSR conditions: hPSCs were differentiated into SNs as described previously^17^. In E8/6 conditions: Differentiations were done as previously described^13^. Briefly, prior to differentiation, plates were coated with vitronectin (5 μg/mL) and incubated for 1h at RT. hPSCs were harvested using 0.5 mM EDTA, 3.08 M NaCl in PBS for 20 min and plated at a density of 200,000 cells/cm^2^. On day of plating (day 0) and day 1, cells were fed with NC differentiation media (day 0-1) containing: Essential 6 Medium, 10 μM SB431542, 1 ng/mL BMP4, 300 nM CHIR99021, and 10 μM Y-27632. BMP4 concentration is very sensitive at this stage and was titrated for each line. Accordingly, BMP4 was not used with iPSC-FD-S3 cells. From day 2 to 12, cells were fed every other day with NC differentiation media (day 2-12): Essential 6 Medium, 10 μM SB431542 0.75 μM CHIR99021, 2.5 μM SU5402, and 2.5 μM DAPT.

On day 12, cells were replated at a density of 250,000 cells/cm^2^ onto plates coated with 15 μg/ml poly-L-ornithine, 2 μg/ml laminin-1, and 2 μg/ml human fibronectin (PO/LM/FN). Cells were incubated with Accutase for 20 min, washed with PBS, and resuspended in SN Media: Neurobasal media containing N2, B-27, 2 mM L-glutamine, 20 ng/ml GDNF, 20 ng/ml BDNF, 25 ng/ml NGF, 600 ng/ml of laminin-1, 600 ng/ml fibronectin, 1 μM DAPT and 0.125 μM retinoic acid. Cells were fed every 2-3 days until day 20. On day 20, DAPT was removed. Differentiation progress was followed using a brightfield microscope (Leica). Genipin (Sigma or Biomaterials), 1,10 anhydrogenipin (AG), dimethyl pimelimidate (DMP, ThermoFisher), bis-(Sulfosuccinimidyl) suberate (BS3, CovaChem), N-hydroxysulfosuccinimide (Sulfo-NHS, ThermoFisher) were added on day 2 of the differentiation and included every media change. Genipin and AG were resuspended in DMSO and aliquoted. DMP, BS3, and sulfo-NHS were resuspended in DMSO (DMP and BS3) or PBS (sulfo-NHS) immediately prior to use.

### Sympathetic neuron (symN) differentiation

The detailed differentiation protocol was described in previous publications^17, 75^. Briefly, hPSCs were dissociated using EDTA and replated on Geltrex (Invitrogen, A1413202)-coated plates at 125×10^3^/cm^2^ density in Essential 6 medium+0.4 ng/ml BMP4 (PeproTech, 314-BP)+10 μM SB431542 (R&D Systems, 1614)+300 nM CHIR99021 (R&D Systems, 4423). On day two, cells were fed with Essential 6 medium+10 μM SB431542+0.75 μM CHIR99021 every two days until day 10 for neural crest induction. Day 10 neural crests were dissociated using Accutase and replated on ultra-low attachment plates (Corning, 07 200) in Neurobasal medium+B27+L-Glutamine+3 μM CHIR99021+ 10 ng/ml FGF2 to induce sympathoblast spheroids. On day 14, spheroids were dissociated using Accutase and replated on PO/LM/FN-coated plates at 1×10^5^/cm^2^ density in Neurobasal medium+B27+L-Glutamine+25 ng/ml GDNF+25 ng/ml BDNF+25 ng/ml NGF+200 μM ascorbic acid (Sigma, A8960)+0.2 mM dbcAMP (Sigma, D0627)+0.125 μM retinoic acid. Medium was changed every three days from day 14 to 20. After day 20, neurons were fed weekly until desired timepoints.

### Prefrontal cortical neuron (PFC) differentiation

The detailed differentiation protocol was described in previous publications^55, 76^. Briefly, hPSCs were dissociated using EDTA and replated on Matrigel (Corning, 356234)-coated plates at 260,000 cells/cm^2^ density in Essential 8 medium supplied with Y-27632. Next day (defined as day 0), cells were fed with E6+100nM LDN193189 (Selleck Chemicals, S2618)+10 µM SB431542 +2 µM XAV939 (TOCRIS, 3748). On day 2, cells were fed with E6 + 100 nM LDN193189 + 10 µM SB every two days until day 8. On day 8, cells were dissociated using Accutase and replated as droplets on PO/LM/FN-coated plates (cells from each well of the 6-well plate were replated into 10 droplets, 10 μl per droplet) in Neurobasal+N2+B27 (1:1000)+FGF8 (50 ng/mL, R&D)+SHH (25 ng/mL, R&D). On day 16, droplets were dissociated using Accutase and replated on PO/LM/FN-coated plates at 1×10^6^ cells/cm^2^ density in Neurobasal+N2+B27 (1:1000)+FGF8 (50 ng/mL). On day 22, neurons were dissociated using Accutase and replated on PO/LM/FN-coated plates at 100,000 cells/cm^2^ density in Neurobasal+N2+B27 (1:50)+FGF8 (50 ng/mL). Neurons were fed without FGF8 after day 40.

### Electrophysiology

Experiments were performed using a Maestro Pro (Axion Biosystems) MEA system. BioCircuit MEA 96 plates containing 8 embedded electrodes/well were coated with Poly-L-ornithine/laminin/fibronectin (as previously described), seeded with day 12 NCCs (250,000 cell/cm^2^) and allowed to continue differentiating. Repeated recordings were made every 2-3 days at 37°C with a sampling frequency of 12.5 kHz for 5 min. Recordings from at least 6 wells per reading were averaged. Firing frequency was normalized to the number of active electrodes.

### Survival and neurite growth assays

Day 12 NCCs, cells were replated on 4-well plates, at 250,000 cells/cm^2^ (for survival assay) or 25,000 cells/cm^2^ (neurite growth assay), coated with PO/FN in SN media with 1 ng/ml NGF. Media was changed every 2-3 days. DAPT was removed after day 20. Cells were fixed on day 13, 20, 27, and 34 and stained for BRN3A and TUJ1 (survival assay) or on day 25 (neurite growth assay) and stained for TUJ1.

### Axotomy models

*For SN regeneration*, day 25 SNs were dissociated with Accutase for 45min. Cells were then collected in a conical tube and filled 1X with PBS. Cells were then centrifuged and resuspended in SN media with DMSO or genipin (10 µM). SNs were seed in PO/LM/FN-coated plates at 250,000 cells/cm^2^ and incubated for 6 days. RNA was collected on the indicated days. Alternatively, SNs were fixed as previously described at the indicated times.

*For PFC regeneration*, day 40 PFCs were dissociated using Accutase and replated on PO/LM/FN-coated plates at 25×10^3^/cm^2^ density with DMSO or genipin (10 µM). Replated neurons were fed every two days until day 45 and fixed for evaluation.

*For symN regeneration*, day 14 sympathoblasts were replated on the PO/LM/FN-coated microfluidic culture devices (eNUVIO, OMEGA4) on one side at 100,000 cells/cm^2^ density. The replating side was defined as the cell body chamber, while another was defined as the axon chamber. NGF containing medium was given to both chambers. On day 20, NGF was given only to the axon chamber to induce axon outgrowth for 10 days. On day 30, axons in the axon chamber were removed using the suction, and DMSO or genipin containing media were added to both chambers for 48 hours. On day 32, regenerating axons were fixed for evaluation.

### Antibodies

The following primary antibodies were used: SOX10 (Santa Cruz, cat# sc-365692), TFAP2A (Abcam, cat# ab108311), BRN3A (Millipore, cat# MAB1585), TUJ1 (Biolegend, cat# 801201), ISL1 (DSHB, cat# 39.4D5-c), PRPH (Santa Cruz, cat# sc-377093), Actin (BD Biosciences, cat# 612656), YAP1 (Proteintech, cat# 13584-1-AP), Phalloidin-iFluor 488 (abcam, cat# ab176753). The following secondary antibodies were used: From ThermoFisher: goat anti-mouse IgG1 AF488 (cat# A21121), goat anti-mouse IgG2a (cat# A-21131), goat anti-mouse IgG2b (cat# A21242), donkey anti-rabbit AF647 (cat# A31573), donkey anti-mouse AF488 (cat# A21202), goat anti-mouse HRP (cat# 62-6520), and goat anti-rabbit HRP (cat# 65-6120). The dilutions used are indicated in each section.

### Flow cytometry

Cells were dissociated with Accutase for 30 min and then washed in Flow buffer (DMEM, 2% FBS, and 1mM L-glutamine). Cells were centrifuged at 200 g for 4 min, resuspended in cold PBS, counted, and diluted to a concentration of 1×10^6^ cells/100μL. Cells were then centrifuged and resuspended in 300 μL BD Cytofix/Cytoperm (BD Biosciences) buffer and incubated on ice for 30 min. Cells were centrifuged for 4 min at 2,000 RPM and resuspended in 600 μL of cold BD Permeabilization/Wash buffer (BD Biosciences). 30 μL goat serum was added followed by incubation on ice for 30 min. Cells were divided in 3: unstained control, secondary antibody control, and sample to stain (200 μL each). All tubes were centrifuged for 4 min at 2,000 RPM and the cells were resuspended in 200 μL of Antibody buffer: BD perm/wash buffer + 10 μL goat serum with or without BRN3A antibody (1:100). Cells were incubated o/n at 4°C. Cells were then washed twice with 300 μL BD Permeabilization/Wash buffer, resuspended in Antibody buffer with or without AF488 goat-anti-mouse (1:500), and incubated on ice for 30 min. Cells were then washed 3X with BD perm/wash buffer. Cells were filtered and analyzed using a Cytoflex S (Beckman). Analysis was done using FlowJo.

### Immunoblotting

Whole cell lysates were obtained from day 12 NCCs or day 20 SNs. Cells were washed once with PBS and incubated with RIPA buffer (ThermoFisher) with 1 mM PMSF and Phospho-STOP (Roche) for 15min on ice. Cells were then vortexed for 10 s and centrifuged at 12,000 RPM for 10 min at 4°C. Protein concentration from supernatants was measured and ran in 7.5% polyacrylamide gels using MOPS buffer (GenScript) at 130 V. Proteins were transferred to a nitrocellulose membrane and blocked for 30 min in 5% non-fat dry milk in 0.1% Tween-20 in TBS (TBS-T, 50 mM Tris-HCl, 150 mM NaCl, pH7.6). Primary antibodies were added to blocking buffer (SOX10 - 1:1000, BRN3A - 1:1000, and Actin - 1:5000) and membranes were incubated overnight at 4°C. Blots were then washed 3X with 0.1% TBS-T and incubated with goat anti-mouse HRP antibody (1:5000) for 1 h at room temperature. Blots were washed 3X with 0.1% TBS-T followed by incubation with Clarity Western ECL Substrate (BioRad). Chemiluminescence signal was detected using UVP ChemStudio (Analytic Jena).

### Immunofluorescence

NCCs and SNs (day 12 and day 20, respectively, unless indicated otherwise) from either 24- or 4-well plates were fixed with 4% paraformaldehyde for 20 min at RT. Cells were washed with PBS and incubated for 20 min with Permeabilization buffer containing 1% BSA, 0.3% Triton-X, 3% goat or donkey serum and 0.01% sodium azide in PBS. Cells were then incubated with the indicated primary antibodies (SOX10 – 1:100, TFAP2A – 1:500, BRN3A – 1:100, TUJ1 – 1:1500, ISL1 – 1:200, PRPH – 1:100) in Antibody buffer containing 1% BSA, 3% goat or donkey serum and 0.01% sodium azide overnight at 4°C. The cells were washed 3X in PBS and incubated for 1h with Secondary antibodies in Antibody buffer. Cells were washed with PBS and incubated with DAPI (1:1,000) for 5min, washed with PBS and stored at 4°C. All imaging was done using a Lionheart FX fluorescence microscope (BioTek). All image analysis and quantifications were done in Fiji. For quantifications, 5 different fields were imaged and quantified. To measure the area of NCCs, day 12 NCCs images were analyzed using Gen5. A mask measuring DAPI signal intensity was used. Threshold was established as the 25% signal intensity from the average signal of each field. The areas of objects between 100 µm to 1000 µm where DAPI signal above background were measured. To measure neurite length, images were transformed to 8 bit grayscale images and neurites were measured using NeuronJ ^77^. For confocal microscopy, on day 12, 50,000 NCCs were seeded in iBidi dishes (cat# 80426) in the presence or absence of genipin. On day 20, SNs were fixed and steined as previously described. Primary antibodies used: TUJ1 – 1:1500, YAP1 – 1:100. Phalloidin-iFluor 488 (1:1000) was incubated with secondary antibodies for 1 h. Imaging was done in an Olympus FV1200 Confocal Laser Scanning Microscope using Argon and Helium-Neon lasers. Images were taken as Z-stacks of 3 µm of height. ImageJ was used to obtain maximum intensity projections and to measure the signal intensity profiles.

### RT-qPCR

RNA was isolated using Trizol (ambion) according to manufacturer’s conditions, resuspended in 20 μL RNase-free water and concentration was measured using NanoDrop One (Thermo Scientific). 1 μg of RNA was converted to cDNA using iScript cDNA Synthesis kit (BioRad) according to manufacturer’s instructions and diluted 1:100 in water. RT-qPCR reactions were run with 1 ul of cDNA and SYBR Green Supermix (BioRad) according to the manufacturer’s conditions. Reactions were run in a C1000 Touch Thermal Cycler CFX96 (BioRad) using the following cycling parameters: 95°C for 5 min, 40 cycles of 95°C for 5s and 60°C for 10 s. Results were analyzed using the comparative CT method. GAPDH was used as a housekeeping gene.

### Primers

The following primers were used in this study: SOX10f-5’CCAGGCCCACTACAAGAGC, SOX10r-5’CTCTGGCCTGAGGGGTGC, TFAP2Af-5’GACCTCTCGATCCACTCCTTAC, TFAP2Ar-5’GAGACGGCATTGCTGTTGGACT, FOXS1f-5’ATCCGCCACAACCTGTCACTCA, FOXS1r-5’GTAGGAAGCTGCCGTGCTCAAA, P75f-5’CCTCATCCCTGTCTATTGCTCC, P75r-5’GTTGGCTCCTTGCTTGTTCTGC, NGN1f-5’GCCTCCGAAGACTTCACCTACC, NGN1r-5’GGAAAGTAACAGTGTCTACAAAGG NGN2f-5’CAAGCTCACCAAGATCGAGACC, NGN2r 5’AGCAACACTGCCTCGGAGAAGA, BRN3Af-5’AGTACCCGTCGCTGCACTCCA, BRN3Ar-5’TTGCCCTGGGACACGGCGATG, ISL1f-5’GCAGAGTGACATAGATCAGCCTG, ISL1r-5’GCCTCAATAGGACTGGCTACCA, PRPHf-5’GTGCCCGTCCATTCTTTTGC, PRPHr-5’GTGCCCGTCCATTCTTTTGC, TRKAf-5’CACTAACAGCACATCTGGAGACC, TRKAr-5’TGAGCACAAGGAGCAGCGTAGA, TRKBf-5’ACAGTCAGCTCAAGCCAGACAC, TRKBr-5’GTCCTGCTCAGGACAGAGGTTA, TRKCf-5’CCGACACTGTGGTCATTGGCAT, TRKCr-5’CAGTTCTCGCTTCAGCACGATG, YAP1f-5’ TGTCCCAGATGAACGTCACAGC, YAP1r-5’ TGGTGGCTGTTTCACTGGAGCA, CYR61f-5’ GGAAAAGGCAGCTCACTGAAGC, CYR61r-5’ GGAGATACCAGTTCCACAGGTC, SULF1f-5’ GGTCCAAGTGTAGAACCAGGATC, SULF1r-5’ GACAGACTTGCCGTCCACATCA, S100A8f-5’ ATGCCGTCTACAGGGATGACCT, S100A8r-5’ AGAATGAGGAACTCCTGGAAGTTA, S100A14f-5’ CCTCATCAAGAACTTTCACCAGTA, S100A14r-5’ GGTTGGCAATTTTCTCTTCCAGG, COL12A1f-5’ CAGTGCCTGTAGTCAGCCTGAA, COL12A1r-5’ GGTCTTGTTGGCTCTGTGTCCT, TMEM2f-5’ GGAATAGGACTGACCTTTGCCAG, TMEM2r-5’ TTCTGACCACCCTGAAAGCCGT, ATF3f-5’ CGCTGGAATCAGTCACTGTCAG, ATF3r-5’ CTTGTTTCGGCACTTTGCAGCTG, SPRR1Af-5’ GTGAAACAACCTTGCCAGCCTC, SPRR1Ar-5’ TGGCAGGGCTCTGGAACCTTG

### RNAseq

#### Global gene clustering

For the first round, paired-end RNA sequencing was performed on hPSC-ctr-H9, iPSC-ctr-C1, and iPSC-FD-S2 day 15 SNs (differentiated as previously described ^17^) with no replicates. Reads were mapped to the human genome build hg19 using TopHat software ^78^. Reads that aligned with no more than 2-bp mismatches were accepted for downstream analysis. After mapping, we computed the expression count matrix from the mapped reads using HTSeq (https://htseq.readthedocs.io/en/master/) and one of several possible gene model databases. The generated count matrix was then processed using the R Bioconductor package DESeq (http://www.huber.embl.de/users/anders/DESeq/) to normalize the full data set and analyze differential expression between samples.

#### GOterm analysis

Genes that showed significant downregulation in iPSC-FD-S2 SNs vs hPSC-ctr-H9 and iPSC-ctr-C1 SNs were subjected to GO term analysis using the DAVID functional annotation tool (https://david.ncifcrf.gov/tools.jsp). The occurrence of GO terms with a FDR score <0.05 were graphed.

### Synthesis of 1,10-anhydrogenipin

Genipin (1.52 g, 6.7 mmol) and triphenylphosphine (1.85 g, 7.1 mmol) were dissolved in 33.5 mL of distilled methylene chloride, brought to 0°C, and stirred for 15 minutes. To this solution was added diisopropyl azodicarboxylate (DIAD, 1.38 ml, 7.1 mmol) dropwise over 10 min and the mixture was stirred on ice for one hour. The reaction was removed from the ice bath and allowed to be stirred overnight at room temperature. TLC (Rf=0.55 in 5:1 hexane/EtOAc) analysis revealed the presence of the product which was purified by flash chromatography (5:1 hexane/EtOAc) to yield a white solid (0.45g, 33%). Mass spectrometry and NMR analysis matched previously published data ^24^.

### Atomic Force Microscopy measurements

To prepare the samples for AFM imaging and Young’s Modulus measurements, Day 25 sensory neurons (SNs) were differentiated in the presence of DMSO or genipin in Nunclon Delta surface dishes (ThermoFisher, cat# 150460 vendor). Cells were washed with PBS and fixed with 4% paraformaldehyde for 20 min at room temperature, followed by 3X washes with PBS. Alternatively, SN-derived extracellular matrix (ECM) was isolated as previously described^79^. SNs were washed with PBS and stripped by incubation with 20mM Ammonium Hydroxide with agitation for 5min. The dishes were then washed 5X with water and the ECM was fixed with 4% paraformaldehyde for 20 min at room temperature followed by 3 washes with PBS. AFM measurements were done using an Agilent 5500 SPM system. Before the AFM experiments, the spring constant of the conical AFM tip on a rectangular cantilever (NANOSENSORS qp-BioAC) was calibrated to be 6pN/nm. The sensitivity of the cantilever was obtained using a very hard glass substrate to be 34.15nm/V. The AFM tip was roughly aligned on the surface of the cell by using an optical microscope, then contact AFM mode was applied to obtain the high-resolution topographic images of the samples. Large area (∼80µm^2^) AFM images were obtained. For each sample, we chose multiple spots on the sample to measure the force-distance curves and calculate the YMs so that a reliable YM can be statically determined. Force-distance curves were further fitted with Hertz model ^46^ (suitable for fitting under small deformation in nanoindentation measurements) and calculate the YM of individual cell by a commercial software PUNIAS ^80, 81^. We carefully control the applied force (setpoint) of 60pN to minimize the deformation. ECM was isolated as previousy described ^79^. Briefly, on day 20, SNs were washed with PBS and incubated for 5 min with 20 mM Ammonium Hydroxide (Sigma, cat# 221228-100ML-A) with constant shaking. Dishes were washed with 5 mL of de-ionized H_2_O and fixed with 4% PFA for 20 min. Cells were then washed 3 times with PBS prior to processing for AFM.

### Animal experiments

*Mice. Elp1^flox/+^and Elp1^Δ^*^20^*^/+^*mice (previously named Ikbkap^9^) were housed and bred at the University of Tennessee Health Science Center Comparative Medicine Department animal core facility.

#### Timed-pregnancies and Genipin treatment

Female *Elp1^flox/+^*mice were crossed with *Elp1^Δ^*^20^*^/+^* males ^9^ for timed-pregnancies. At E10.5 a fresh mouse cage was provided to pregnant female mice for acclimatization and shortly after breeder mouse chow was replaced with genipin-containing moist breeder chow (500 mg/Kg of chow), corresponding to daily consumption of 75 mg of genipin per Kg of body weight. Genipin-containing moist chow was supplied fresh daily. Females were weighed daily and sacrificed at late gestation (E18.5).

#### Genotyping of embryos

Genomic DNA was prepared from embryo tail biopsies, and embryos were genotyped by PCR amplification as described previously ^9^.

#### Histological analyses

Embryos were fixed in 4% paraformaldehyde in phosphate-buffered saline (PBS) for at least one week. Embryos and placentas were weighed and then incubated for 24 hours at 4°C in PBS containing 0.25 M sucrose, 0.2 M glycine; dehydrated; cleared with toluene; and embedded in paraffin. Paraffin blocks were serially sectioned at 7 μm, mounted in superfrost slides (Fisher) and stained with haematoxylin and eosin (H&E).

#### Volumetric determination and neuronal counts

Volumes of sensory ganglia were determined as described previously^9, 29^. In brief, H&E-stained serial paraffin sections (7 μm) spanning the whole ganglia were analyzed under a Zeiss stereomicroscope, and width and length were measured every 5^th^ section. Volumetric measurements were performed by calculating and adding the volumes between every section analyzed. For neuronal counts, neurons with clearly visible nucleoli were counted from photomicrographs of H&E-stained paraffin sections. Total neuronal numbers were estimated based on the total volume of the ganglia.

#### Immunohistochemistry on paraffin sections

For immunohistochemistry on paraffin sections (IHC-P), slides were deparaffinized, rehydrated, and incubated with 0.3% H_2_0_2_ in methanol for 20 min, to quench endogenous peroxidase. Sections were then washed with PBS, blocked for 1 hr with 4% BSA, 0.2% Triton X-100 in PBS, and incubated at 4°C for 24hrs with primary rabbit monoclonal anti-CGRP antibody (Calbiochem PC205) diluted 1:200 in 0.4% BSA; 0.2% Triton X-100 in PBS. After several washes in PBS, primary antibody detection was carried out using the Vector ABC Elite kit according to manufacturers’ instructions, followed by incubation with DAB brown substrate (BD Biosciences).

#### Quantification of CGRP+ sensory nociceptive neurons

Nociceptive neurons were quantified at E18.5 by counting CGRP-positive staining of neurons in cross sections of lumbar L1 DRG. For each embryo, four images per DRG from at least two independently-immunostained sections were captured at 20X and divided into quadrants. Total number of cells and number of CGRP-positive cells per quadrant were determined using ImageJ. Positive signal was established as 30% above background.

### Statistical analysis

All analysis and graphs were done using PRISM (GraphPad). Number of independent experiments (biological replicates, n) and statistical analysis are indicated on each figure legends. Biological replicates are defined as independent differentiations started at least 3 days apart or from a freshly thawed vial of cells. Statistics for animal data, see below. Data presented are shown as mean ± SD. For experiments *in vivo:* data were derived from multiple independent experiments from distinct mice. Animal studies were performed without blinding of the investigator. Histological analyses were performed in at least 3 embryos per genotype and treatment. No statistical method was used to predetermine sample size, but sample size was based on preliminary data and previous publications. In all experiments the differences were considered significant when p<0.05. The differences between groups were assessed using one-way ANOVA followed by Tuckey HSD post hoc test.

### Study approval

Animal experiments were carried out in strict accordance with the Guide for the Care and Use of Laboratory Animals of the National Institutes of Health. The protocol was approved by the Animal Care and Use Committee of the University of Tennessee Health Science Center.

## List of Supplementary Materials

Sup Fig. 1 to Sup. Fig. 6

Sup. Table 1 to Sup. Table 3

## Acknowledgements

We thank Michael Tiemeyer and Yao Yao for critical discussions. Harold ‘Skip’ Ralph at the Weill Cornell Microscopy and Image Analysis Core facility for image analysis. Julie Nelson from the UGA Cytometry Shared Resource Laboratory for her help with flow cytometry experiments.

## Author Contributions

KS-D conceived, conducted and analyzed experiments, and wrote manuscript. PD and ID conducted animal experiments and tissue analysis. AJP conducted experiments. CW, ARP, and GB synthesized 1,10-anhydrogenipin, SC provided the chemical library, LS provided support, advice, and funds for the chemical screen. NZ conceived, designed, led the study, conducted experiments, wrote the manuscript and provided funds.

## Data and materials availability

All data needed to evaluate the conclusions in the paper are present in the paper or the Supplementary Materials. Requests for reagents should be directed to the corresponding author, Nadja Zeltner

---

**Supplementary Figure 1.**
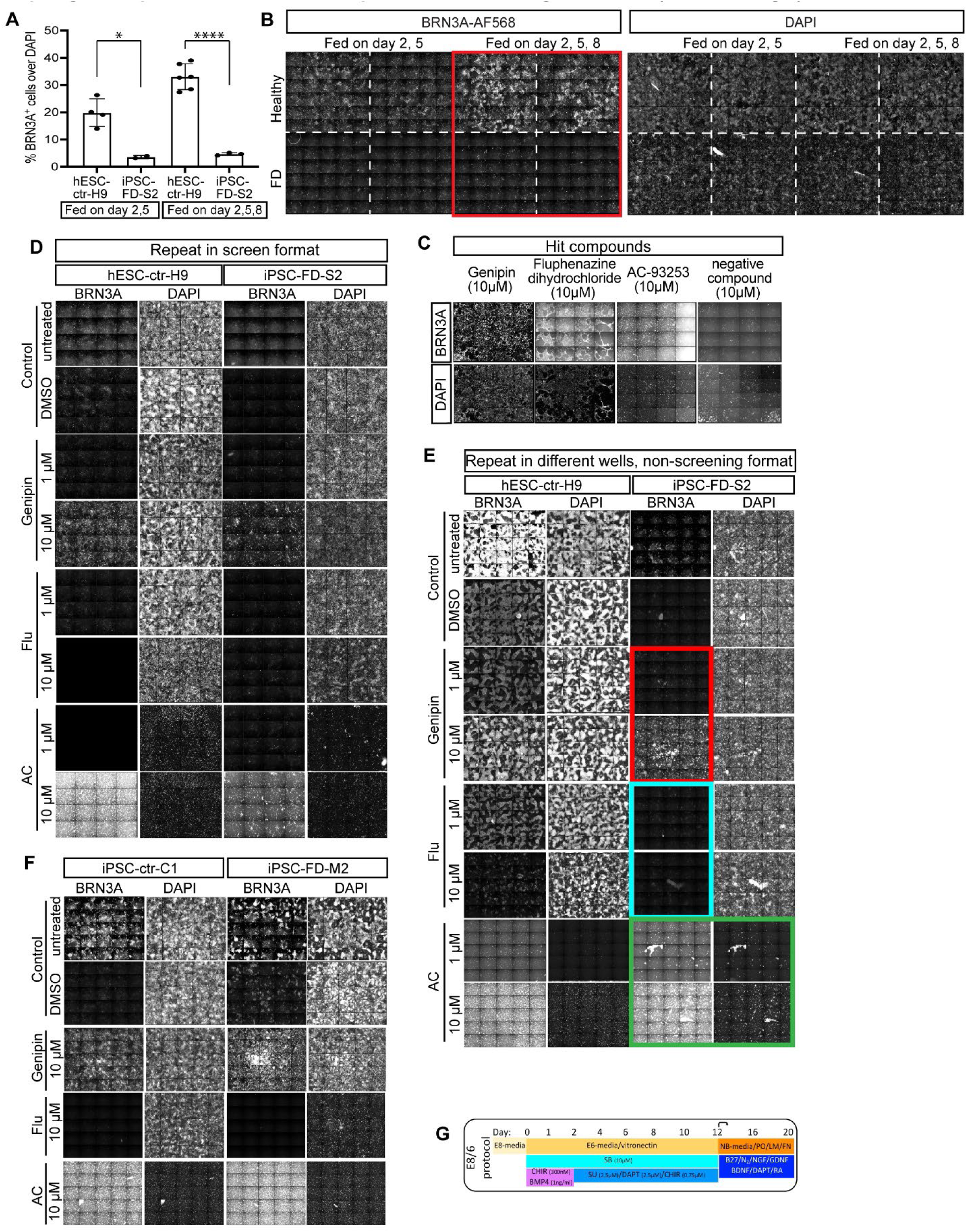
Adaptation of the SN differentiation protocol to high-throughput screening conditions and hit validation. **A)** Quantification of the number of BRN3A^+^ SNs over total DAPI^+^ cells. Two-tailed Student’s t-test, *p<0.01. **B)** Imaging of 30 fields per 96 well for each condition. **C)** Example wells are shown from the screen. Repeat of the SN differentiation in the presence of the hit compounds in 96 wells in screening conditions outlined in Fig. 1C. Whole wells are shown, imaged in 16 tiles. **E)** Repeat of SN differentiation in the presence of the hit compounds in a different well format, i.e. 48 wells and using non-screening conditions outlined in Fig. 1A. Whole wells are shown, imaged in 25 tiles. **F)** Effect of hit compounds tested on additional cell lines, i.e. healthy iPSCs and FD iPSCs from patients with mild FD symptoms (iPSC-FD-M2). Non-screening differentiation conditions were used in a 48 well format. All differentiations were performed in KSR conditions. **G)** Schematic representation of the SN differentiation in E8/E6 conditions (as previously reported^1^).

**Supplementary Figure 2.**
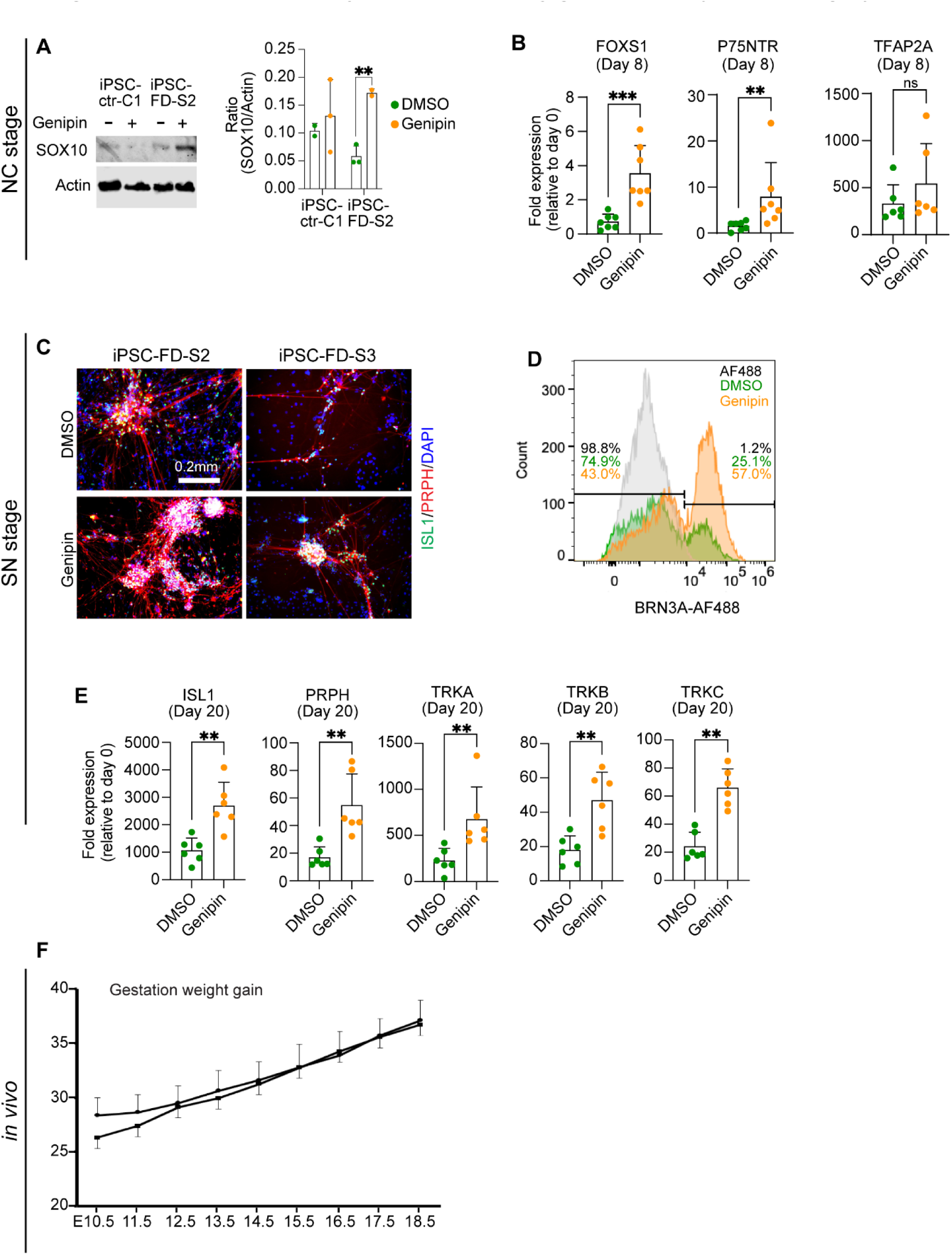
*in vitro* and *in vivo* developmental phenotypes in FD are rescued by genipin (related to Fig.3 and Fig. 4). **A)** Immunoblotting of SOX10 from day 12 NC cells. Representative blot is shown (left) and quantification (right, n=2-3 biological replicates). **B)** RT-qPCR-based gene expression analysis of SN-primed NC at day 8 (n=6-7 biological replicates). **C)** Genipin restores SN differentiation. Day 20 SNs were treated with genipin and fixed and stained using the indicated antibodies (n=5 biological replicates). **D)** Representative histogram of BRN3A signal measured by flow cytometry. **E)** RT-qPCR-based gene expression analysis of SNs at day 20 (n=6 biological replicates). **F)** Gestational female weight gain in genipin-treated (squares, n=6) and untreated (circles, n=3) dams. For **A, B** and **D**, Two-tailed Student’s t-test. ns, non-significant, **p<0.005, ***p<0.001. All graphs show mean ± s.d. For **B** and **D**, iPSC-FD-S2 and iPSC-FD-S3 data are pooled as FD.

**Supplementary Figure 3.**
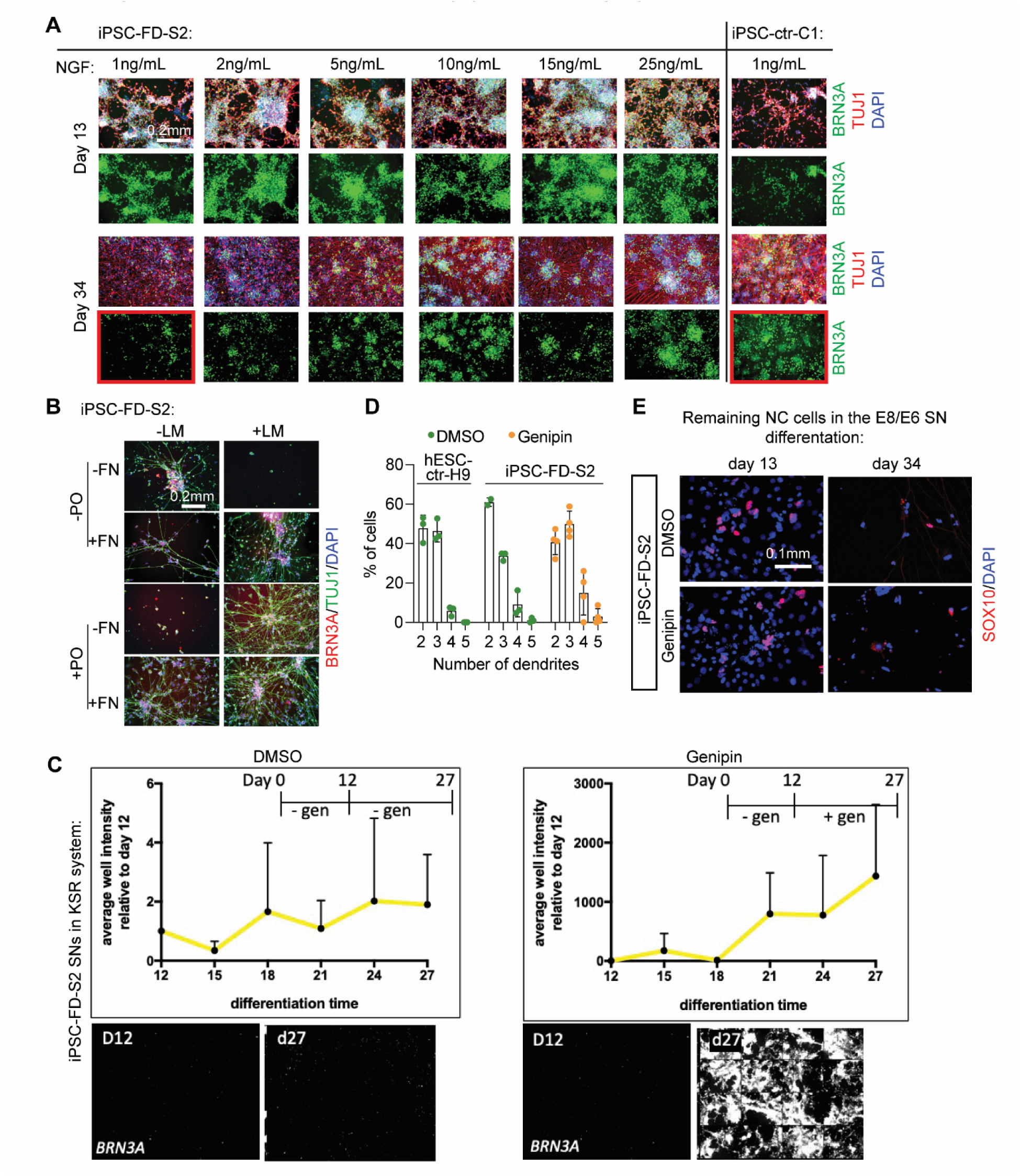
SN survival assay adaptation and characterization (related to Fig 5). **A)** NGF titration to assess SN survival over 21 days by IF in iPSC-FD-S2 and healthy iPSC-ctr-C1 cells. **B)** Exclusion experiment to assess which of the surface coating proteins LM, PO, FN are required in iPSC-FD-S2 SNs. **C)** Survival assay performed in KSR conditions (Fig 1A). 16-tile images are shown of SNs at day 12 and day 27, with or without genipin treatment (bottom). Quantification of BRN3A signal intensity is plotted relative to day 12 (top). **D)** Number of neurites of hPSC-ctr-H9 and iPSC-FD-S2 SNs differentiated in the presence of genipin (10 µM) were quantified. n=2-4 biological replicates. **E)** Few SOX10+ NC cells are present in the SN differentiation at day 13 and 20. All graphs show mean ± s.d.

**Supplementary Figure 4.**
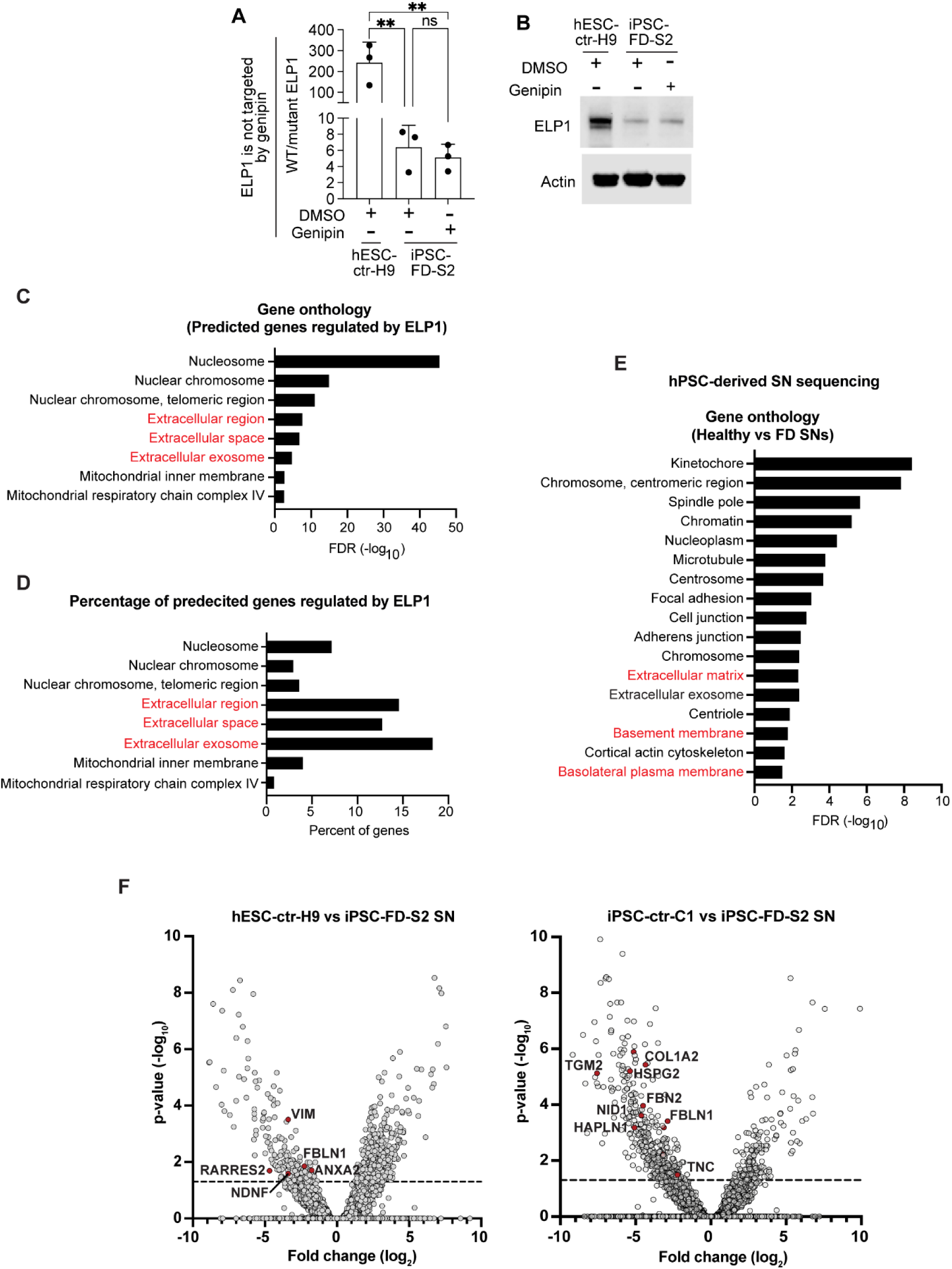
Effects of genipin in ELP1 and expression of ECM-related genes in FD mouse and iPSC-derived FD SNs. **A)** Treatment with genipin of NC cells up to day 8 does not alter *ELP1* splicing inefficiency (RT-qPCR). n=3 biological replicates. one-way ANOVA followed by Tukey’s multiple comparisons. ns, non-significant, **p<0.005. **B)** Genipin does not change ELP1 protein levels in FD (immunoblot). n=3 biological replicates. Representative blot is shown. **C,D)** Analysis of genes predicted to be controlled by Elp1 in FD mouse. **C)** Gene ontology of predicted genes that are regulated by Elp1. **D)** Percentage of genes predicted to be regulated by Elp1. **E, F)** RNA sequencing in hPSC-derived SNs comparing healthy (hPSC-ctr-H9 and iPSC-ctr-C1) versus FD (iPSC-FD-S2). E) Gene ontology analysis of significantly downregulated genes in FD vs healthy SNs. **F)** Differential gene expression of hPSC-ctr-H9 vs FD SNs (left) and iPSC-ctr-C1 vs FD SNs (right). ECM-related genes downregulated in FD are highlighted. Dotted line indicates significance threshold (p<0.05).

**Supplementary Figure 5.**
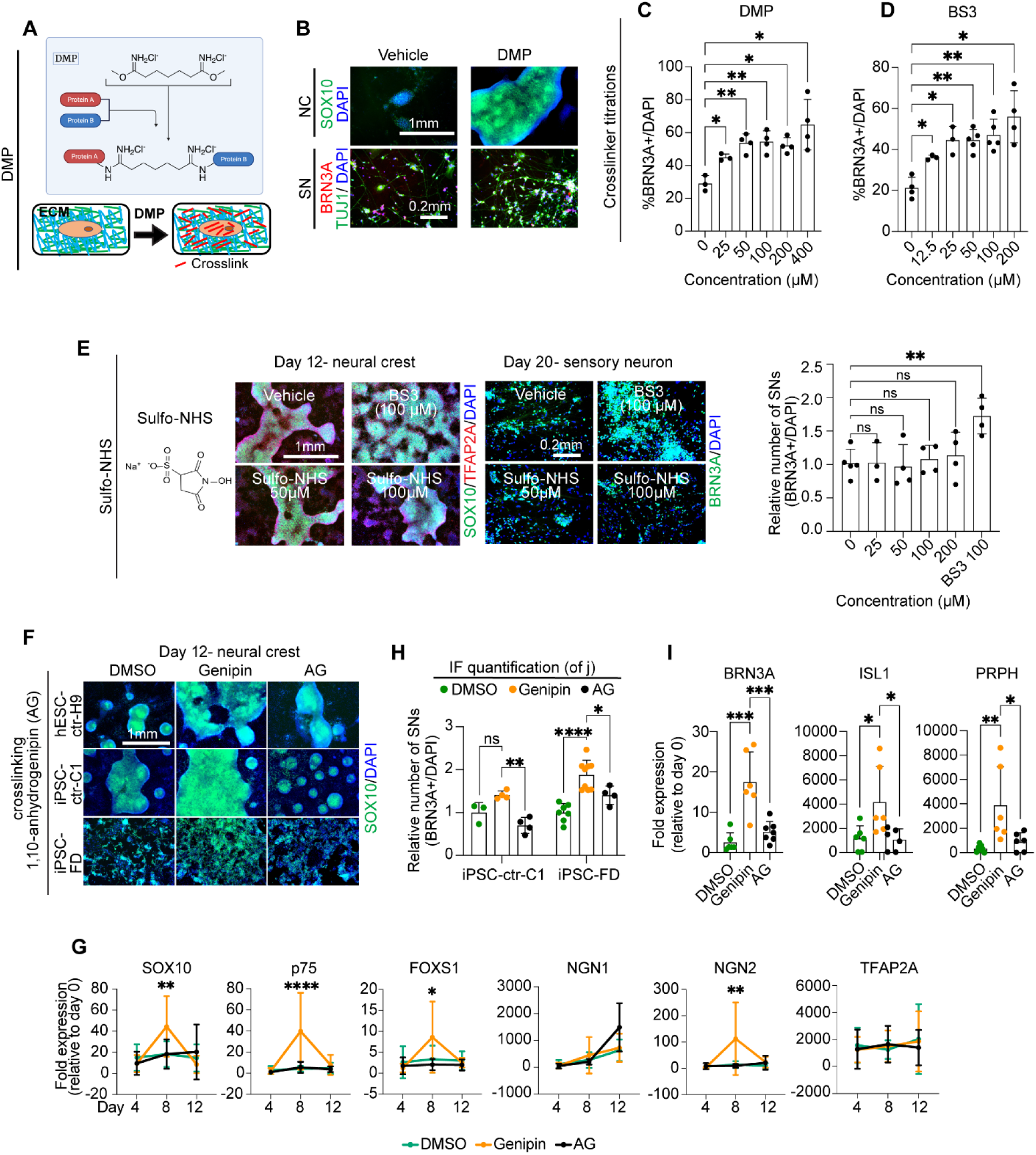
Genipin’s mode of action is through crosslinking of ECM proteins (related to Fig 6). **A)** Schematic of DMP crosslinking action and its intracellular/extracellular location. **D)** DMP rescues the NC and SN differentiation defect in FD. iPSC-FD-S2 cells were differentiated in the presence of DMP and fixed on day 12(NC) and day 20 (SN). Following staining using the indicated antibodies. n=3-4 biological replicates. **C)** DMP titration on iPSC-FD-S2-derived SNs on day 20. n=3-4 biological replicates. one-way ANOVA followed by Tukey’s multiple comparisons. *p<0.05, **p<0.005. **D)** BS3 titration on iPSC-FD-S2-derived SNs on day 20. n=3-5 biological replicates. one-way ANOVA followed by Tukey’s multiple comparisons. *p<0.05, **p<0.005. **E)** Sulfo-NHS (inactive BS3) does not rescue NC formation, nor SN formation in iPSC-FD-S2 cells. n=3-5 biological replicates. one-way ANOVA followed by Tukey’s multiple comparisons. ns, non-significant, **p<0.005. **F)** Genipin that has been chemically altered to delete its crosslinking effects (1,10-anhydrogenipin) is not capable to rescue NC formation in FD cells. **G-I)** 1,10-anhydrogenipin does not properly restore NC cells (**G**) or SNs (**H** and **I**) compared to genipin. Assessed by RT-qPCR (G and I) and IF quantification (**H**, related to Fig. 6H). For **G**, n=7-8 biological replicates. Two-way ANOVA followed by Šídák multiple comparisons. For **H** and **I**, n=6-8 biological replicates. one-way ANOVA followed by Tukey’s multiple comparisons. *p<0.05, **p<0.005, ***p<0.001, ****p<0.0001. All graphs show mean ± s.d. For **F, G, H,** and **I,** iPSC-FD-S2 and iPSC-FD-S3 data are pooled as FD.

**Supplemental Figure 6.**
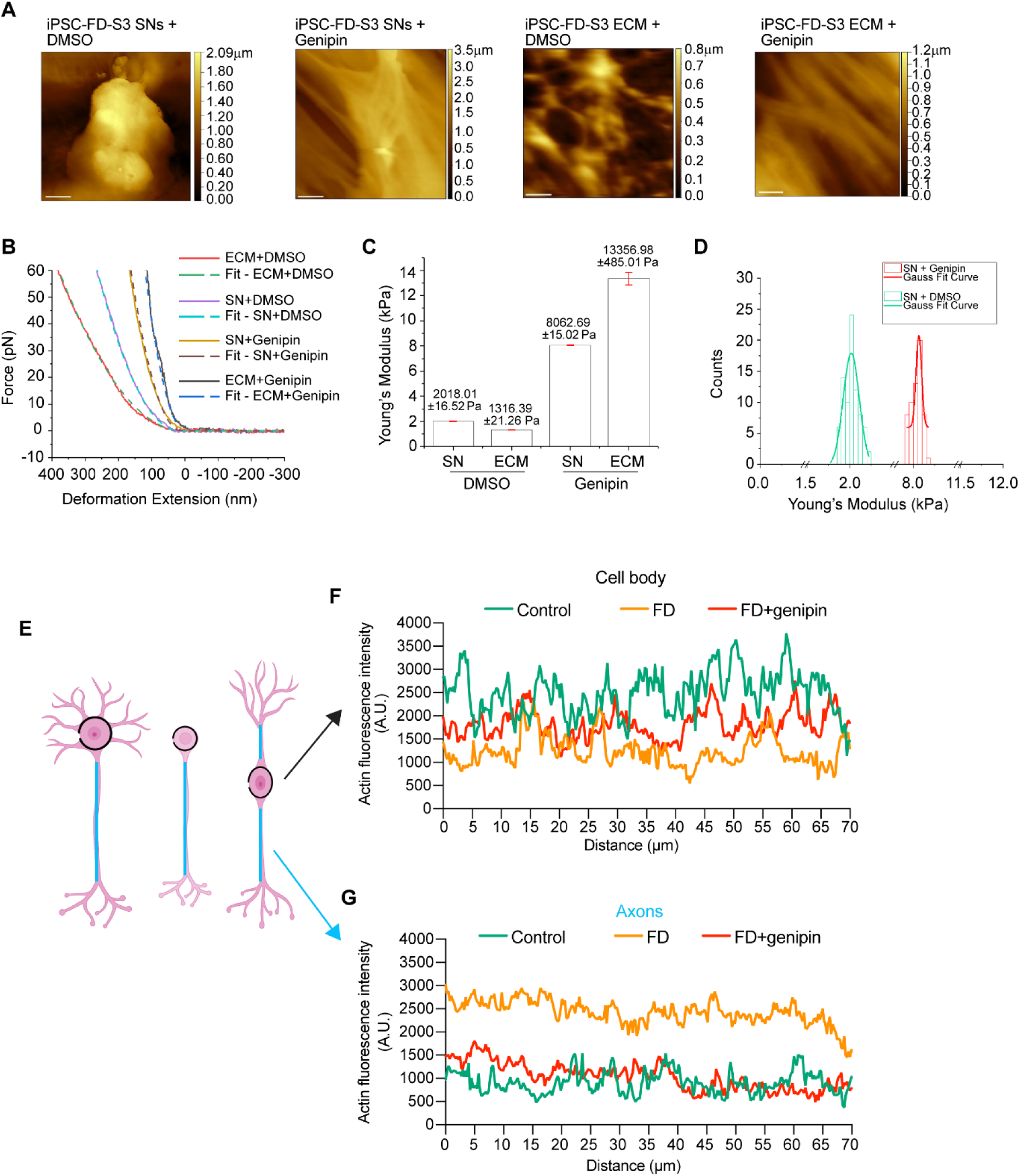
Effects of Genipin crosslinking of ECM (related to Fig 6). **A)** AFM topographic images (Scale bar: 10µm). **B)** Force-distance curves (solid) recorded by force spectroscopy, along with Hertz model fitted curves (dash) which was used to calculate the Young’s Modulus. **C)** Comparison of Young’s Modulus for four different samples from a representative experiment. **D)** Histogram of Young’s Modulus for SNs in DMSO (green) and genipin (red) with Gaussian distribution. A Gaussian distribution fit curve is overlayed to highlight sample variability. **E)** Schematics for actin signal measurement in cell bodies (black) and axons (blue) in SNs with different *in vitro* morphology. **F, G)** Quantification of actin signal intensity from images in **Fig.6K** (n=7-10 cells from 3 biological replicates).

**Supplementary Table 1.** Small molecule screening data

| Category | Parameter | Description |
| --- | --- | --- |
| Assay | Type of assay | IF staining |
|  | Target | Increase in sensory neuron differentiation |
|  | Primary measurement | IF staining of BRN3A+ cells |
|  | Key reagents | Differentiated sensory neurons |
|  | Assay protocol | Adapted from Chambers et al., 2012 |
|  | Additional comments |  |
| Library | Library size | 640 compounds, i.e. half of the LOPAC library |
|  | Library composition | Cell signaling and neuroscience |
|  | Source | Sigma |
|  | Additional comments |  |
| Screen | Format | 96 well plates |
|  | Concentration(s) tested | 1mM and 10mM |
|  | Plate controls | DMSO treated wells, healthy hPSCs (+ control) and disease cells without compounds (- control) |
|  | Reagent/ compound dispensing system | Manual multi-pipettor |
|  | Detection instrument and software | MetaXPress software: Cell scoring Module from Molecular Devices |
|  | Assay validation/QC | + and – controls and DMSO only wells (see above) |
|  | Correction factors | n/a |
|  | Normalization | DAPI+ cells and DMSO only treated wells |
|  | Additional comments |  |
| Post-HTS analysis | Hit criteria | Phenotype reproducible in screening and non-screening format |
|  | Hit rate | 1 |
|  | Additional assay(s) | Phenotype reproducible in different well format<br>Phenotype reproducible in additional control and disease hPSC lines |
|  | Confirmation of hit purity and structure | n/a |
|  | Additional comments |  |

**Supplementary Table 2.** Hit compounds

| Compound name | Screen [c] | Fold change over DMSO (average) |
| --- | --- | --- |
| DMSO only | 10µM | 4.5 |
| hPSC-ctr-H9 in DMSO | 10µM | 7.7 |
| iPSC-FD in DMSO | 10µM | 1.6 |
| Fluphenazine dihydrochloride | 10µM | 17.5 |
| AC-93253 iodide | 1µM | 16.5 |
| genipin | 1µM | 14.2 |
| genipin | 10µM | 16.1 |

**Supplementary Table 3.** Genipin does not affect normal embryonic development

|  | Total embryos | Live embryos | Resorptions | Number of litters |
| --- | --- | --- | --- | --- |
| No treatment | 46 | 40 | 6 | 6 |
| + Genipin | 44 | 41 | 3 | 6 |

